# A study on experimental bias in post-translational modification predictors

**DOI:** 10.1101/2022.11.28.518163

**Authors:** Jasper Zuallaert, Pathmanaban Ramasamy, Robbin Bouwmeester, Nico Callewaert, Sven Degroeve

## Abstract

Protein post-translational modifications (PTMs) play an important role in numerous biological processes by significantly affecting protein structure and dynamics. Effective computational methods that provide a sequence-based prediction of PTM sites are desirable to guide functional experiments. Whereas these methods typically train neural networks on one-hot encoded amino acid sequences, protein language models carry higher-level pattern information that may improve sequence based prediction performance and hence constitute the current edge of the field. In this study, we first evaluate the training of convolutional neural networks on top of various protein language models for sequence based PTM prediction. Our results show substantial prediction accuracy improvements for various PTMs with current procedures of dataset compilation and model performance evaluation. We then used model interpretation methods to study what these advanced models actually base their learning on. Importantly for the entire field of PTM site predictors trained on proteomics-derived data, our model interpretation and transferability experiments reveal that the current approach to compile training datasets based on proteomics data leads to an artefactual protease-specific training bias that is exploited by the prediction models. This results in an overly optimistic estimation of prediction accuracy, an important caveat in the application of advanced machine learning approaches to PTM prediction based on proteomics data. We suggest a partial solution to reduce this data bias by implementing negative sample filtering, only allowing candidate PTM sites in matched peptides that are present in the experimental metadata.

**Availability and implementation:** The prediction tool, with training and evaluation code, trained models, datasets, and predictions for various PTMs are available at https://github.com/jasperzuallaert/PhosphoLingo.

**Supplementary information:** Supplementary materials are available at *bioRxiv*.

## Introduction

Proteome-wide analysis of post-translational modifications (PTMs) is currently accomplished through liquid chromatography-mass spectrometry (LC-MS) experiments^1–3^ that require a protease to digest proteins into peptides for further analysis. Recent advancements in open modification searching methodologies for LC-MS data analysis enabled the high-throughput identification of numerous types of PTMs within a given sample^4^. However, this approach is constrained by inherent limitations, as only a subset of the proteome and its associated PTMs is observable via LC-MS. Notably, peptides falling below or exceeding the MS detection range cannot be detected, and numerous peptides lack sufficient proteotypic characteristics or abundance for reliable detection and identification. This gap can be closed computationally through *in-silico* models that generalize the LC-MS data, predicting potential modification sites that have yet to be experimentally observed. Consequently, this computational strategy facilitates downstream MS data interpretation and provides guidance for subsequent *in-vitro* and *in-vivo* experiments^5^.

The most commonly applied protease is trypsin, cleaving proteins after arginine and lysine residues, resulting in a positive charge at the peptide C-terminal side-chain that is advantageous for MS and MS/MS analysis. Alternative proteases that cleave at other residues^1^ are applied to increase the proportion of a proteome that can be analysed using LC-MS.

One of the most frequently studied PTMs is phosphorylation^6^, where the availability of large data repositories has resulted in an extensive range of sequence-based prediction models. Consequently, our study on performance enhancement and methodological bias analysis of machine learning approaches is focused on phosphorylation site (from here onward referred to as phosphosite) prediction, while confirming the resulting conclusions for acetylation, methylation, sumoylation, and ubiquitination. In protein phosphorylation, phosphate groups are transferred by enzymes from adenosine triphosphate to specific residues in a protein sequence, with serine, threonine or tyrosine residues being the most common targets in eukaryotes. Kinases and phosphatases are involved in virtually all biological processes by targeting specific sites for modification. The substrate preferences of kinases depend on the structural and physicochemical properties of the target substrate. Phosphorylation sites (from here onward referred to as phosphosites) are most often found at accessible and disordered regions of a protein^7^. In addition, the amino acid sequence and structure at the phosphosite are important, with for example proline-directed kinases predominantly targeting S/T residues followed by a proline residue^8^.

The majority of phosphosite prediction models utilize neighbouring sequence data to predict which serine (S), threonine (T) or tyrosine (Y) residues are likely to be phosphorylated^5,9–27^. In addition to general phosphosite prediction, some approaches also include kinase family-specific or kinase-specific predictions^9,12,13,15,17,18,21–23,25^. Over the last two decades, different machine learning algorithms have been applied, such as random forests^10,14,24^, support vector machines^9,16,24^, and gradient boosting trees^9^. These classical approaches require numerical feature vector representations engineered from the amino acids surrounding a candidate phosphosite, which we will refer to as the receptive field. Manually crafted input features for phosphorylation prediction models include physicochemical properties^14,16,21,26^, information theory features^15,17,28^, as well as additional information such as structural features (e.g., secondary structure)^10,16,24^, functional features^10,24^, and protein-protein interactions^17,25^.

Deep learning based model architectures such as convolutional neural networks (CNNs)^13,17– 20,22,26^ and long short-term memory networks (LSTMs)^20,26^ have been shown to advance phosphosite prediction. These models learn low-level feature representations from the sequence input data during training, alleviating the need for preliminary manual feature engineering. Recently, LMPhosSite utilized protein language models (pLMs) to improve phosphosite prediction by adding single-position embeddings as input features^28^. PLMs are pretrained models that yield enriched, structure-aware sequence representations, instead of merely encoding the amino acid composition of a receptive field in a protein^29–35^. They have demonstrated value in various tasks, such as few-shot contact map prediction^36^, protein structure prediction^34^, zero-shot mutation impact prediction^37^, or phylogenetic relationship modelling^38^.

In our approach, we utilize various pLM embeddings for entire protein fragments, on top of which we then fit a CNN. We confirm that the enhanced information content contributes substantially to the prediction of accuracy compared to standard one-hot encoding-based representations, as well as to publicly available state-of-the-art tools.

After this, we undertook the core of our study, which is the use of model interpretation methods to understand which patterns are considered relevant by the machine learning models. Our results reveal that, in addition to known local and global sequence patterns, the models exploit LC-MS protease-specific patterns. This results in overly optimistic prediction performance estimations for *de novo* phosphorylation site prediction. To partially restrain this prediction bias and improve data quality, we suggest a negative sample filtering method based on experimental metadata, which we validated on phosphorylation data from individual experiments with different proteases.

## Results

### PLMs substantially improve PTM site prediction accuracy

We evaluated the most widely adopted pLMs for PTM site prediction by training CNNs on top of the extracted embeddings. We compared our new predictors with CNNs trained on traditional one-hot encodings, as well as with current state-of-the-art predictors that use a variety of machine learning approaches. These predictors traditionally use one-hot encoded representations, with a combination of a CNN module and a capsule network module (MusiteDeep^18^), three densely connected CNN blocks (DeepPhos^19^), or a local context module and a global module built from convolutional and LSTM layers, and a squeeze-and-excitation network (DeepPSP^13^). Finally, LMPhosSite^28^ uses a CNN module on a trainable embedding, in combination with a fully-connected neural network built on the pLM representation for the single phosphosite position. For more information, see the Methods section, and their respective publications.

We studied a selection of PTMs based on data abundance, starting with phosphorylation of serine (S), threonine (T) and tyrosine (Y) as the benchmark case. To extend our findings, we then studied acetylation of lysine (K), methylation of arginine (R) and lysine, sumoylation of lysine, and ubiquitination of lysine. Model training and evaluation datasets were compiled using the commonly adapted procedure that labels candidate PTM sites as positive when the site was confidently observed as modified by LC-MS, and as negative otherwise, when found in proteins with at least one positive annotation (see Methods for more details), indicating sufficient abundance of the protein to allow for PTM detection.

Figure 1 shows the phosphosite prediction performance for the different amino acid representations, as well as for the state-of-the-art predictors. To compare between predictors, we consider the area under the precision-recall curve (AUPRC) as the most meaningful performance metric in highly imbalanced datasets (i.e. with many more non-modified sites than modified sites). Detailed results are shown in Suppl. Table 1.

**Figure 1.**
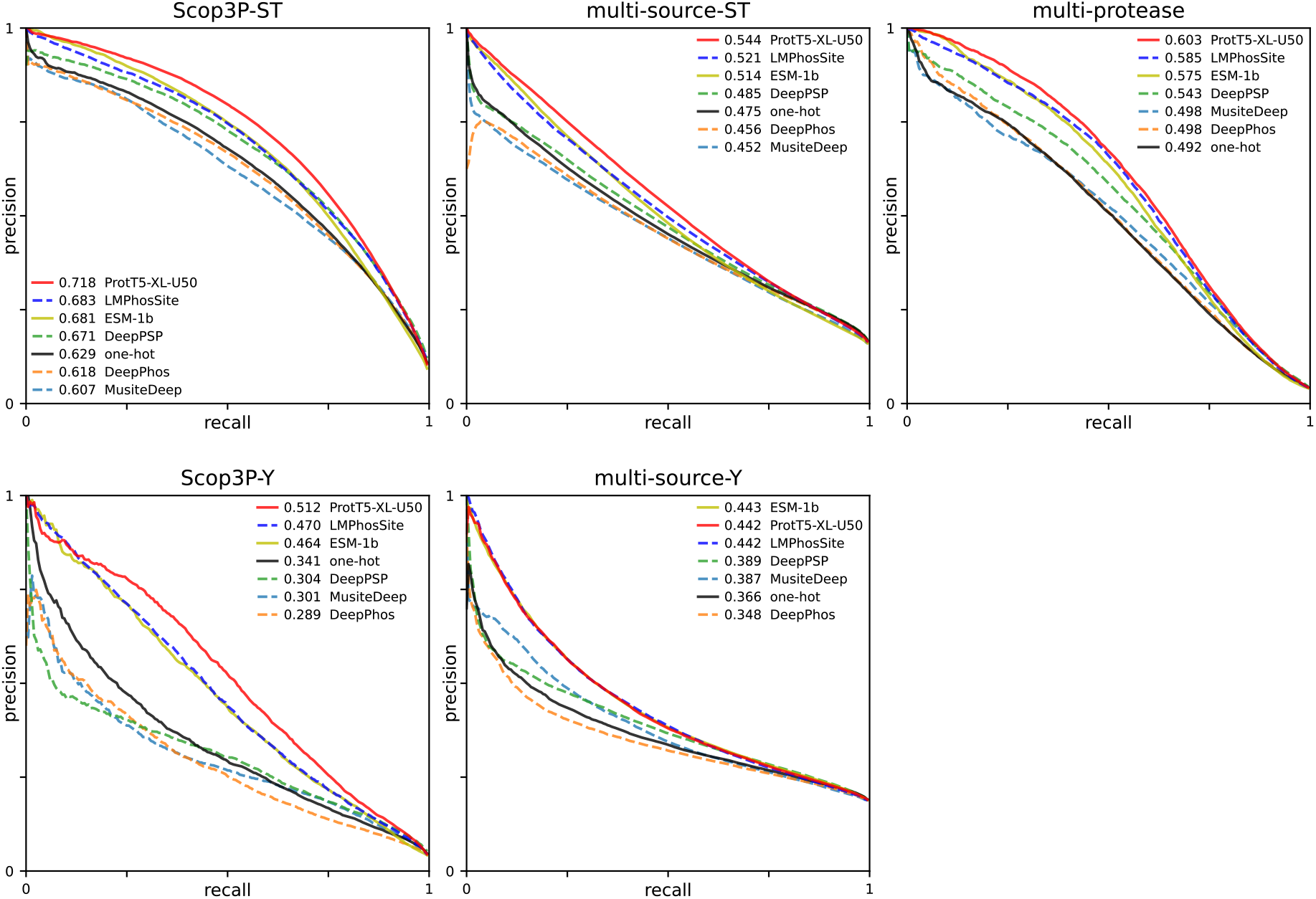
Precision-recall curves for the ProtT5-XL-U50 and ESM-1b pLMs, a one-hot encoding, and existing predictors. Three data sources are considered for phosphosite prediction: high-quality reprocessed Scop3P data^39^, data from multiple traditional sources (UniProtKB/SwissProt, dbPTM^40^, Phospho.ELM^41^, and PhosphoSitePlus^42^) used as the benchmark in other phosphosite prediction publications^13,28^, and combined S/T phosphosites from individual experiments with different proteases^1^. Average curves over 10 runs (per dataset) are depicted. In the legend, the mean AUPRC is reported.

Remarkably, in the evaluation on four out of the five datasets that were considered here, the ProtT5-XL-U50 language model consistently performed best in terms of AUPRC, with improvements of 14.1% to 50.1% over the one-hot encoded CNN, and more modest improvements of 2.0% to 4.6% over the next best pLM-based model. Only on multi-source-Y, ESM-1b performed roughly equally (+0.2%). Furthermore, the ProtT5-XL-U50 outperformed the one-hot encoded state-of-the-art models by 7.0% to 68.4% on all datasets considered. Compared to the LMPhosSite tool that also employs ProtT5-XL-U50 based representations but with a different model, our CNN model showed an improvement of 2.3% to 8.9% in terms of AUPRC, except again on multi-source-Y, where performance was roughly equal.

As shown in Figure 2 and Suppl. Table 2, for the other PTMs we also observe a consistent improvement of the prediction performance when replacing one-hot encoded sequences with ProtT5-XL-U50 based sequences representation, resulting in relative improvements in AUPRC between 13.9% and 37.2%.

**Figure 2.**
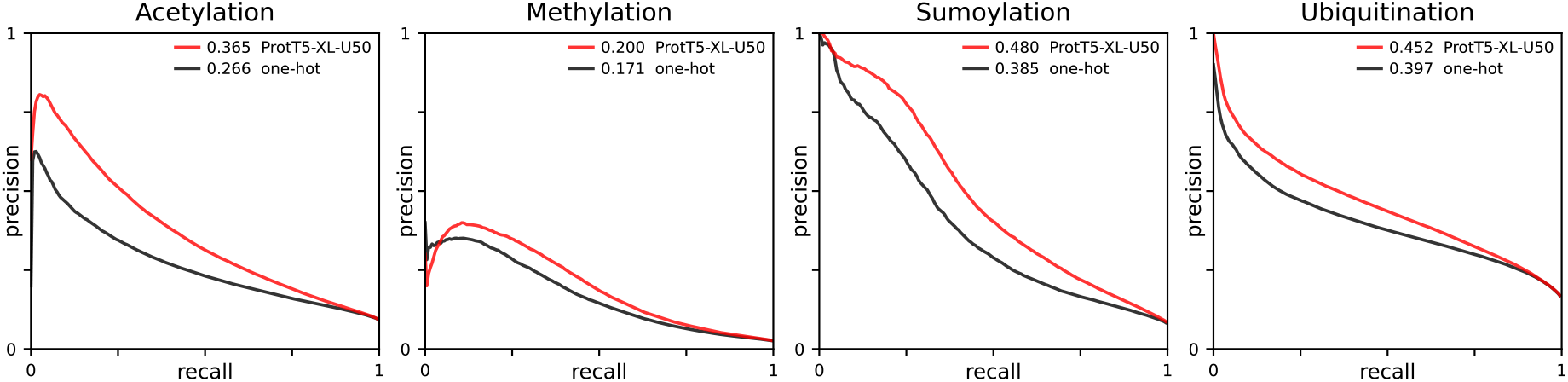
Precision-recall curves for the ProtT5-XL-U50 pLM and a one-hot encoding, on other PTMs. Average curves over 10 runs (per dataset) are depicted. In the legend, the mean AUPRC is reported.

### Feature importance analysis of phosphosite predictors shows discriminative features from different kinases

We computed SHapley Additive exPlanations (SHAP^43^) to estimate the average importance of individual residues in the proximity of candidate phosphosites. Results for models trained on the Scop3P datasets using the ProtT5-XL-U50 model are visualized in Figure 3, and models using a one-hot encoding are visualized in Suppl. Fig. 2.

**Figure 3.**
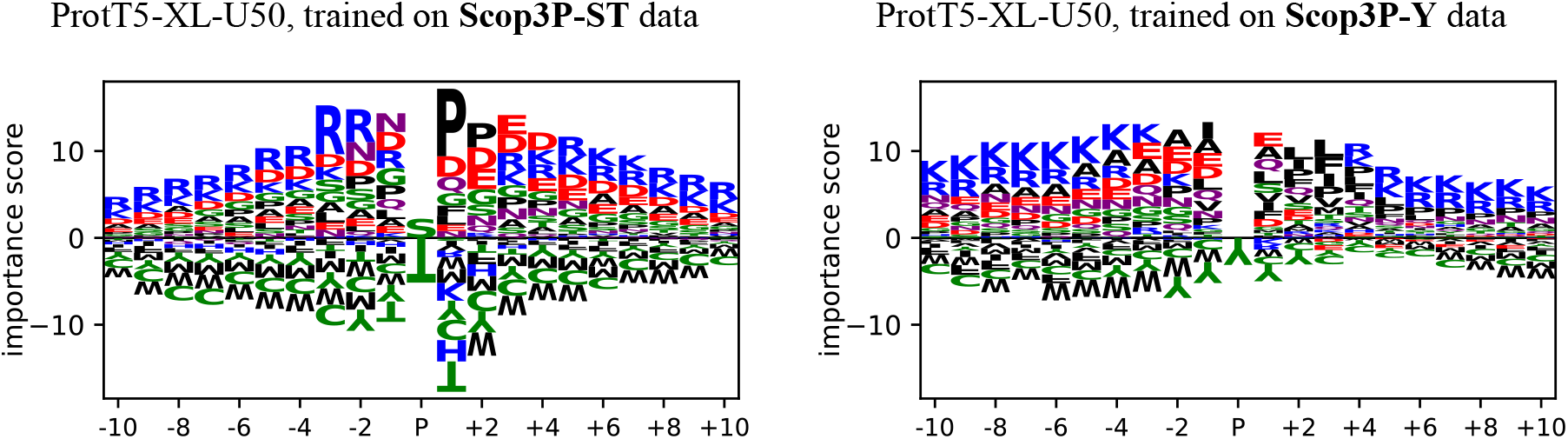
Average importance scores per position calculated using DeepLiftSHAP, for the ProtT5-XL-U50 prediction model setup, on the Scop3P datasets. Visualizations are cropped to show the twenty positions surrounding the candidate phosphosite.

For the ST models, the most notable residue is proline (P) at position P+1, which is known to be a strong signal towards a phosphosite prediction, as several kinases in the CMGC group are known to be proline-directed^8^, such as those belonging to the MAPK and CDK kinase families (Suppl. Fig. 1a-b). Other favourable residues in the ST visualization are arginine (R) at positions P-3 and P-2, as often seen for kinases in the PKa family (Suppl. Fig. 1c), and aspartic(D) and glutamic acid (E) on the first positions downstream of the candidate site, as often seen for kinases in the CK2 family (Suppl. Fig. 1d).

Distinct motifs are also observed when visualizing SHAP values for other PTMs, as shown in Figure 4 for pLMs, and in Suppl. Fig. 3 for one-hot encoded models. For K acetylation, certain amino acids are preferred, like G on positions P-2 and P-1. In contrast to ST phosphorylation, no clear conserved motif is detected here, perhaps due to the diversity of K acetyltransferases and the possible non-enzymatic origin of many acetylation events^44^. Methylation shows a clear preference for G in the proximity of the candidate methylation site, and for sumoylation, the known ΨKXE consensus motif^45^ (with Ψ denoting a hydrophobic residue, and X any residue) can clearly be distinguished in the visualization.

**Figure 4.**
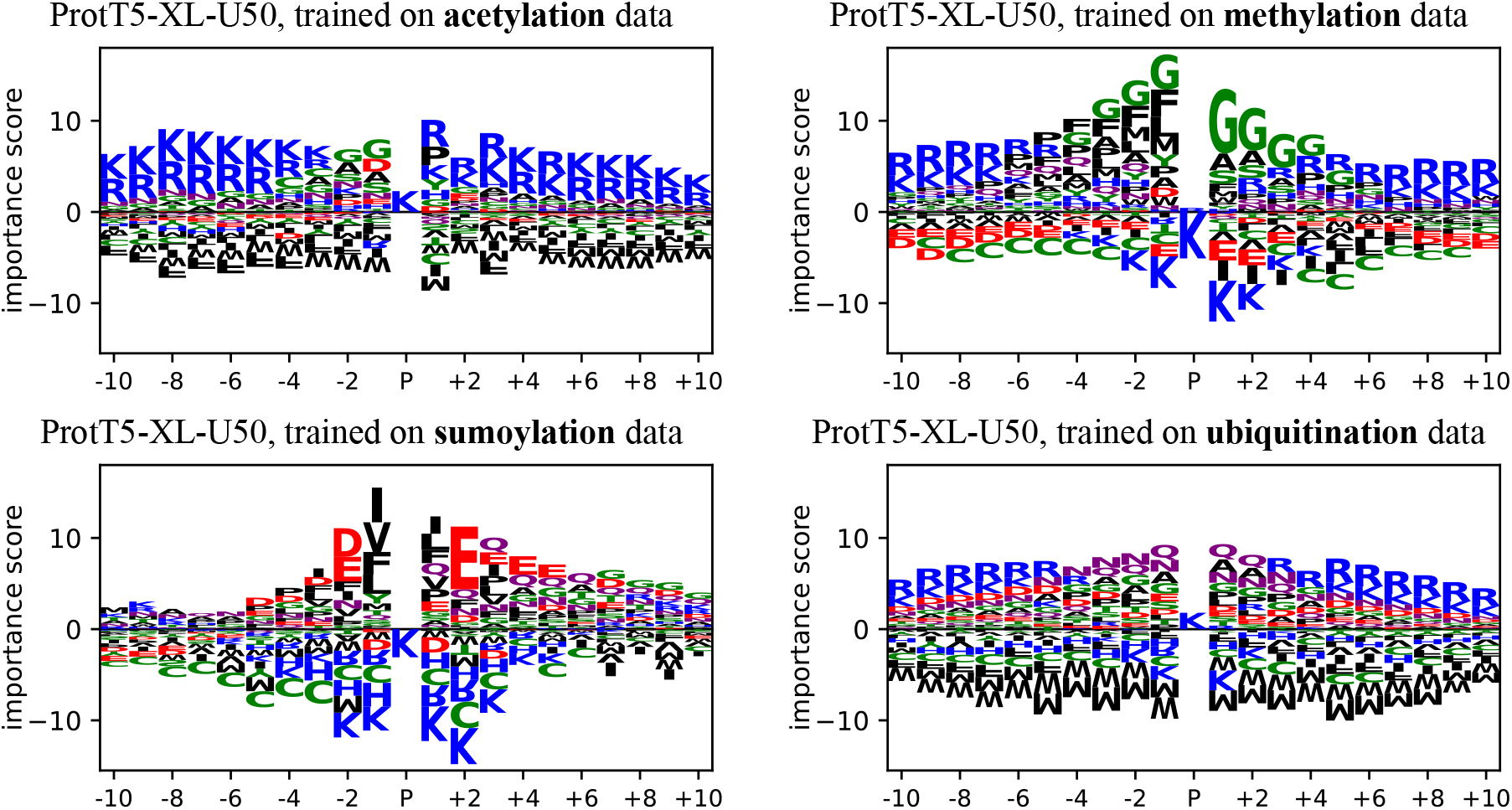
Average importance scores per position, calculated using DeepLiftSHAP, for ProtT5-XL-U50 models trained on the acetylation, methylation, sumoylation, and ubiquitination datasets. Scores are calculated on their respective test sets.

### CNNs trained for PTM site prediction disfavour inaccessible phosphosites

The average importance scores in Figure 3**Figure and** Figure 4 indicate that the models often disfavour hydrophobic amino acids in the proximity of the candidate PTM site. On average, for the pLM-based predictor trained on the Scop3P-ST dataset, W/C/F/Y/M are considered least favourable when in the proximity of a phosphosite. As phosphosites need to be reached by kinases for phosphorylation to take place, the majority of phosphosites are found at the solvent exposed area of the protein (more than 80%, while less than 2% are located in buried regions)^7^. Hydrophobic (A/F/I/L/M/V/Y/W) and C residues are more prevalent in buried regions^46^, suggesting that the models implicitly learn that phosphosites occur less frequently in buried regions. Similar effects with disfavoured hydrophobic residues are observed in tyrosine phosphorylation, acetylation, and ubiquitination, and with disfavoured cysteines in tyrosine phosphorylation, methylation, and sumoylation (Figures 3-4).

This is confirmed by considering prediction distributions for candidate phosphosites in buried, interface and accessible regions. AlphaFold^47,48^ was used to obtain protein structures for all proteins in the dataset that were shorter than 2700 amino acids. Next, DSSP^49^ was used to calculate the solvent accessibility and secondary structure for all candidate phosphosites. Figure 5 shows the distribution of predicted probabilities for positively labelled phosphosites in the evaluation set. Predicted scores of phosphosites located in buried regions tend to be lower than those in interface regions, and much lower than those in accessible regions. Similar conclusions can be drawn from secondary structure analysis, where phosphosites in random coil regions generally score higher than those in helices and sheets, which is in line with statistical analysis of phosphorylation data^7^.

**Figure 5.**
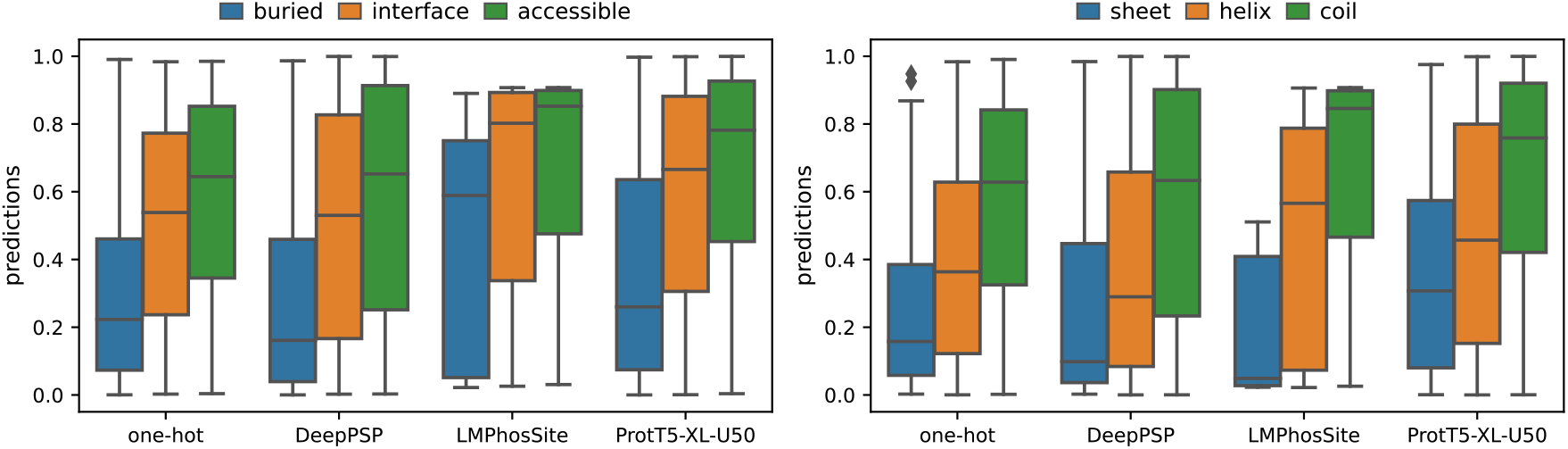
Distribution of predicted probabilities for positively labelled phosphosites in the Scop3P-ST-PF test set, per model, divided by surface accessibility and secondary structure.

### Datasets from experiments with individual proteases show biased learning

Figures 3 and 4 reveal a preference of arginine and lysine residues at all positions beyond a few amino acids upstream of downstream of the candidate PTM sites, except for sumoylation. As the LC-MS data used to compile the training sets were mainly generated using the protease trypsin, this observation strongly points to a protease-induced data bias. We hypothesize that the prediction models learn an enriched presence of tryptic cleavage sites near PTM-sites due to the limits of LC-MS analysis. Indeed, positively labelled datapoints are found in peptides (with an average length of about 10 to 12 amino acids) that contain at least one arginine or lysine, whereas negatively labelled datapoints are obtained from all of the other candidate sites in the protein, including those in long stretches without any arginine or lysine residues.

To test this hypothesis, we performed additional experiments on single-protease datasets, where we trained predictors on phosphorylation data compiled from experiments that applied different proteases. Figure 6 confirms that models trained on single-protease datasets demonstrate consistent enrichment of their corresponding cleavage sites. The amino acids with the highest average importance within the twenty positions surrounding the phosphosites are aspartic acid for AspN, phenylalanine/leucine for chymotrypsin, glutamic acid for GluC, lysine for LysC, and arginine/lysine for trypsin, consistent with the most frequent cleavage sites of the respective proteases. Similar results for one-hot encoded models are shown in Suppl. Fig. 4.

**Figure 6.**
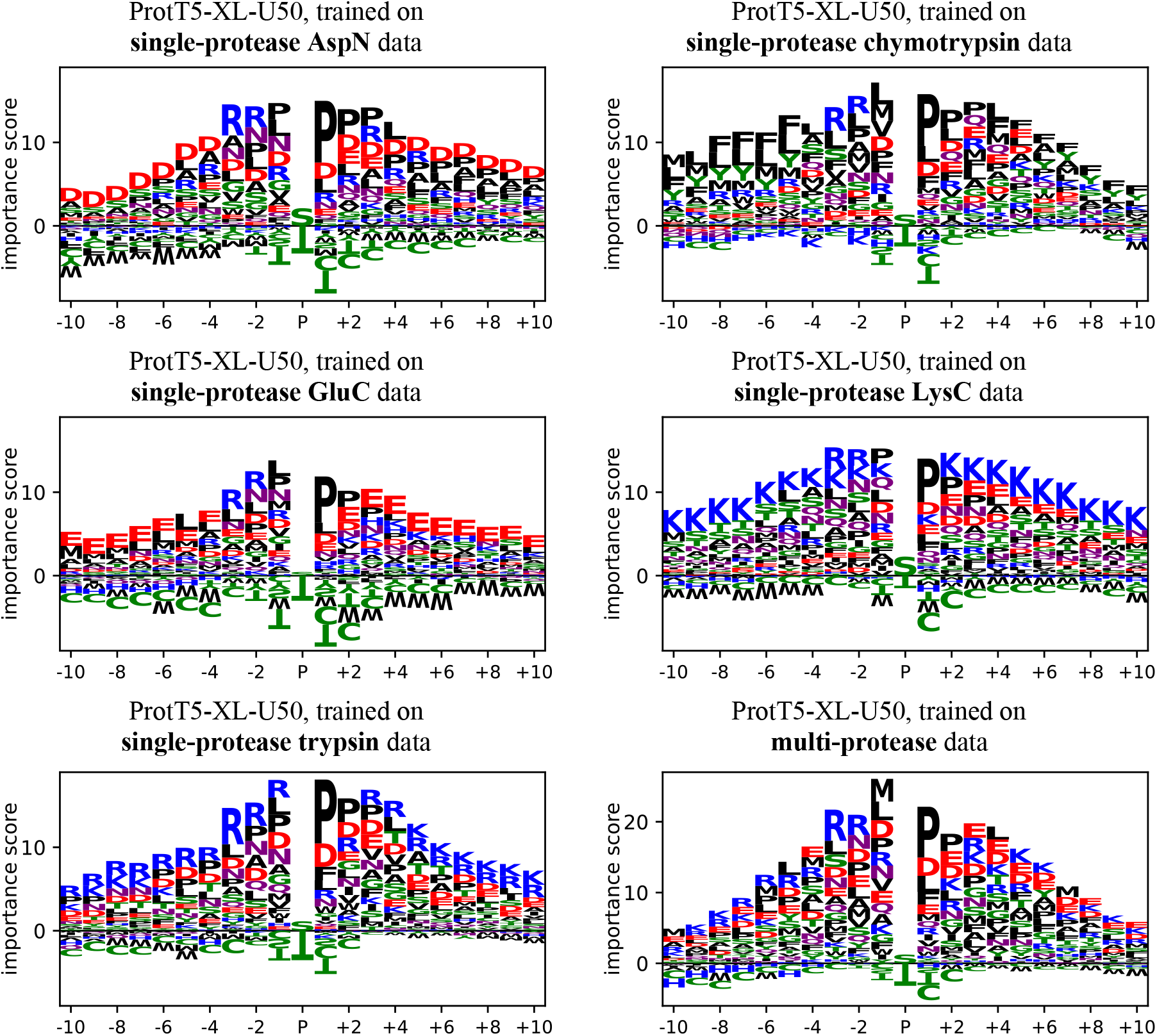
Average importance scores per position, calculated using DeepLiftSHAP, for the ProtT5-XL-U50 prediction model setup, on the single- and multi-protease datasets. Visualizations are cropped to show the twenty positions surrounding the candidate phosphosite

### Peptide filtering rules out non-observable annotations and mitigates protease bias

By default, the “negative” datasets consist of all candidate PTM sites that were not observed as carrying the corresponding PTM in LC-MS experiments, for proteins in which at least one PTM site of the type under study is annotated in the dataset. However, many of these sites will simply not be observable at all in LC-MS proteomics, for instance because they are located in too long peptides, not well-ionizing peptides, etc. Hence, for the datasets where experimental metadata was available, we applied peptide filtering to only retain negative annotations for which the candidate PTM site was actually observed by LC-MS, but never as modified by the PTM. We refer to these datasets as peptide-filtered (PF). In addition to ruling out non-observable (false negative) candidates from training and evaluation, we hypothesized that this would reduce protease-related bias, as prediction models can no longer by default classify all candidate sites far away from any cleavage site (arginine or lysine in the case of trypsin) as negative samples.

We first compared the impact on prediction performance in terms of AUROC, as this metric is not influenced by dataset imbalance as is the case for the AUPRC. In Suppl. Table 3, it is observed that the AUROC drops consistently for all representations by 1.6% to 3.1% when applying peptide filtering on the Scop3P-ST dataset. For Scop3P-Y, a drop in AUROC of 2.4% to 8.3% occurs. This suggests an increased level of complexity for the prediction task, as a subset of easily predictable targets is no longer considered in the evaluation.

Next, the visualizations in Suppl. Figure 5 indeed suggest a slight decrease of the bias levels when applying peptide filtering. We compare the average importance score of R and K residues in the proximity of the phosphosite (disregarding the 7 central amino acids containing other strong signals) to the average importance of the most important residue, P at P+1. For the Scop3P-ST dataset, we observe that the average scores for R and K are 30.0% and 19.5% of the average score for P at P+1, and that this drops to 16.4% and 15.2% respectively when applying peptide filtering. This suggests that the models are less inclined to learn the protease-induced bias. We further verify this in the next section.

### Model performance deteriorates when transferred between individual protease datasets

To further analyse the observed protease-specific bias, we evaluated phosphosite predictors that were trained on different data sources, across different test sets. Concise model transfer results for the pLM-based models are shown in Figure 7, with additional results in Suppl. Fig. 6.

**Figure 7.**
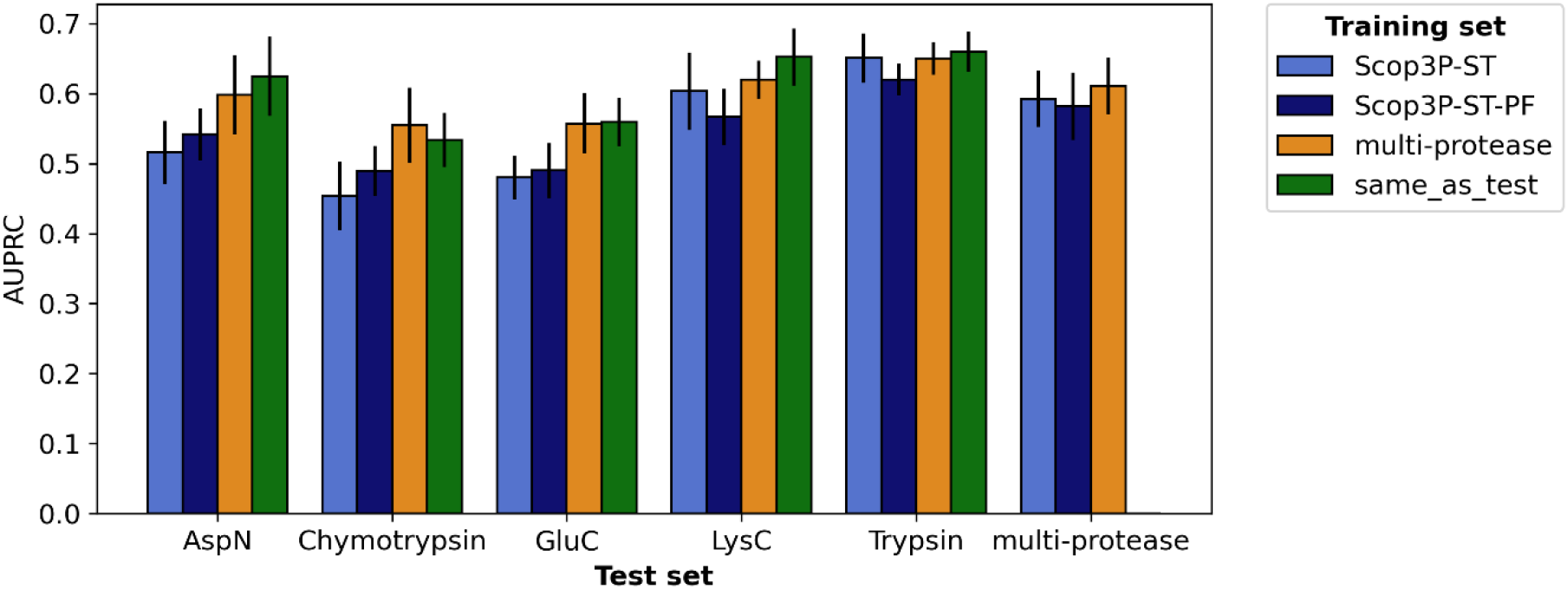
Evaluation on non-peptide-filtered single-protease and multi-protease datasets (x-axis), when training on a different data source (legend). Predictions are computed from ten folds of the evaluation set, where training is performed on either Scop3P-ST data, Scop3P-ST-PF data, multi-protease data, or on the dataset matching the data source on the x-axis. Note that proteins present in the test fold are always omitted from training. The mean AUPRC over these ten test folds and the standard deviations are shown, for ProtT5-XL-U50 models.

ProtT5-XL-U50 models trained on the Scop3P-ST dataset are consistently outperformed by the respective single-protease trained models with a relative AUPRC increase of 16.5% to 21.1%, for the AspN, chymotrypsin, and GluC data. For LysC data, where cleavage occurred at lysine residues (partly sharing the bias with the trypsin-based Scop3P data), the performance difference was smaller (8.1%). For trypsin data, the models trained on Scop3P data almost reached the same level (1.4%), as the Scop3P data is exclusively constructed from experiments using trypsin digestion.

Additionally, we compared the models trained on the Scop3P-ST to models trained on Scop3P-ST-PF data, to investigate the effect of the endeavoured quality improvement in the filtered data. We observe that the trypsin bias learned by the models is reduced, as performance improves on AspN, chymotrypsin, and GluC data by 2.1% to 7.9%. Simultaneously, the performance drops for the LysC and trypsin datasets by 6.0% and 4.7% respectively, as part of the bias that was to the favour of these proteases is reduced.

Models trained on multi-protease data further improve on the Scop3P-ST-PF-based models by 4.9% to 13.8%. Compared to the models trained on single-protease data, the difference in performance is limited, between -4.3% and +3.8%. Overall, this suggests that models trained on data obtained from multiple proteases generalize better towards the proteases included in the data acquisition setup. Extra model transfer results can be found in Suppl. Fig. 6, where we can also see that models trained on the multi-source data generalize worse than the models trained on the quality-enhanced Scop3P datasets.

Finally, we further examined extra filtering steps. In one strategy (Scop3P-ST-PF-MC), we only retained candidate phosphosites from peptides without any missed cleavages, with the intention of balancing out short R/K-rich peptides that are the result of more frequently missed cleavages. In another strategy (Scop3P-ST-PF-Loc), we attempted to remove negative candidate sites that were present in peptides with unlocalized modifications, i.e. where detected phosphorylations could not be unambiguously be assigned to a Ser/Thr residue within the peptides due to low localization scores. However, in both cases the filtering seemed to be less effective, with a drop in performance compared to the peptide-filtered datasets, as clearly illustrated in the additional results in Suppl. Fig. 6. Therefore, we did not implement these approaches in our final strategy.

## Discussion

We examined the potential of pLMs for the prediction of PTM sites from protein sequence data by fitting CNNs on top of protein representations generated by the language models. Our results show a significant improvement in PTM site prediction performance compared to current state-of-the-art neural network architectures that rely on one-hot encoded representations. This improvement was consistent across various phosphorylation datasets obtained from diverse sources, as well as for other modifications, indicating that pLMs offer informative protein sequence representations that substantially improve PTM site prediction accuracy.

Upon inspection of feature importance for serine/threonine phosphorylation prediction, we found evidence supporting known kinase family-specific residue patterns, such as proline at P+1, arginine at P-3 and P-2, and aspartic and glutamic acid at the first positions downstream of the candidate phosphosite. These findings suggest that the proposed models recognize the characteristic features of an ‘average’ phosphosite across all kinases. Further analysis incorporating protein 3-D structure revealed that predicted probabilities were generally higher within accessible regions than within buried regions, and higher within random coil regions compared to those within alpha helices or beta sheets, as is to be expected^7^.

However, our analysis also revealed a strong preference of both existing and our new prediction models towards amino acids that are targeted by proteases for cleavage, particularly in the vicinity of candidate PTM sites.

However, our analysis also revealed that both existing and our new prediction models learn that the nearby presence of the amino acids at which the proteomics protease cleaves increases the probability of a PTM site to be modified. We attribute this pattern to the inherent experimental biases present in bottom-up LC-MS/MS proteomics experiments, rather than being a modification-specific pattern, and this finding is of importance for other machine learning classifiers trained on proteomics datasets.

We hypothesized and validated that this bias arises from multiple sources. First, candidate phosphosites in long stretches of protein sequence lacking cleavage sites cannot be detected in LC-MS/MS experiments. For datasets compiled without applying our proposed peptide filtering, all candidates in those stretches would be labelled as non-phosphosites. Secondly, a higher frequency of missed cleavages near phosphosites^1,3^ results in more arginine- and lysine-enriched peptides that would typically be cleaved into fragments that are too short for MS/MS analysis when not phosphorylated. Thirdly, as also shown in Suppl. Figure 7, localization scores that are calculated to pinpoint the modified residue within a peptide are on average lower for longer peptides, as more candidate residues are present. Hence, the accuracy of phosphosite determination within such longer peptides is lower. A prediction model can exploit these technical biases by utilizing the lack of arginine and lysine around non-phosphosites to steer its prediction to a lower modification probability, and by increasing the phosphosite probability for occurrences within arginine- and lysine-rich regions.

Thus, general phosphorylation prediction models do not just learn biological phosphosite sequence fingerprints to predict the likeliness of phosphorylation, but also the likeliness of detection with current experimental setups. The trypsin-related training bias is also observed in our acetylation, methylation and ubiquitination models.

Further confirming this bias source, phosphosite prediction models trained on datasets obtained using one protease show reduced accuracy when evaluated on datasets obtained using a different protease. Training a model with combined data from individual proteases already showed improved performance when evaluating across the different datasets, and this practice can be recommended for other machine learning work on proteomics data.

To further suppress these bias sources from experimental data acquisition, we investigated a peptide filtering method to improve data quality by removing candidate sites that were never observed in peptides in LC-MS, and for which the annotation as “not modified” was hence based merely on negative experimental data, which has a poor evidence level. In addition to model evaluation on data with fewer false negative data points, models trained on filtered data generalized better when transferring to phosphorylation datasets derived using different proteases for digestion. Hence, such limitation of training data to positively identified observed peptides is good practice. However, the proposed filtering method does not resolve all potential sources of dataset bias, such as the increased frequency of missed cleavages near phosphosites and the lower localization scores for increased peptide length, for which our initial attempts at adding extra filtering steps did not further improve generalizability across proteases.

Potentially, so-called “middle-down” proteomics, in which only limited proteolysis is aimed for, would strongly reduce these biases. Current evolutions in mass spectrometry such as enhanced non-collisional electronic fragmentation may enable the generation of such less biased datasets.

As a final remark, the application for which PTM site predictions are done steers the choice of training data. When doing *de novo* predictions, protease bias is detrimental, and the training data should be as heterogeneous as possible. However, when predicting detectable PTMs for LC-MS/MS data interpretation purposes (e.g., enhanced search engines), a model trained on data acquired using the same protease should be used.

We have released the PhosphoLingo tool that implements the most accurate pLM-based phosphosite predictors presented in this study on our GitHub repository (https://github.com/jasperzuallaert/PhosphoLingo). These models were trained on the Scop3P-ST-PF and Scop3P-Y-PF datasets. We also provide a user-friendly framework to replicate the results reported in this manuscript, and to train and evaluate models on additional phosphorylation datasets, or on other post-translational modifications. Additionally, we provide phosphorylation predictions for all S, T and Y residues in the human proteome, as well as acetylation/sumoylation/ubiquitination predictions for all K residues, and methylation predictions for all R and K residues.

## Methods

### Protein representations

As a baseline, the receptive field was represented as a one-hot encoding of the amino acids, i.e., no pLM was used. Then, multiple pLMs were considered in this study: ESM-1_small_ and ESM-1b^31^, ESM-2^34^, ProtT5-XL-U50^32^, CARP-640M^50^, Ankh-base^35^, and Ankh-large^35^ (see Suppl. Table 4 for more details).

### Neural network architecture

To evaluate the different protein representations for phosphorylation prediction, we constructed a CNN architecture on top of the pLM representations, which we then trained without fine-tuning the pLM further. Different approaches where the representations were further fine-tuned were considered, but ultimately discarded due to risks of overfitting on the huge models, as also suggested in literature^38^. An extensive hyperparameter search was done for each of the initially considered language models (ESM-1_small_, ESM-1b, ProtT5-XL-U50), from which we then constructed a final hyperparameter setup that is used throughout all experiments. We chose to use a single setup that works well for all datasets and representations to increase the robustness and reproducibility of our approach. Details are listed in Suppl. Table 5.

As schematically depicted in Figure 8, the CNN contains a series of convolutional blocks, with each convolutional block consisting of a convolutional layer followed by a regularization layer (either dropout or batch normalization), and finally a max pooling layer. The resulting activations are flattened, after which a fully-connected, a regularization layer, and an output layer are added. Furthermore, all convolutional and fully-connected layers are followed by a rectified linear unit (ReLU), and a sigmoid is added at the end to provide probabilities in output. The default one-hot encoded and pLM-based setups are schematically depicted in Suppl. Figs. 8-9.

**Figure 8.**
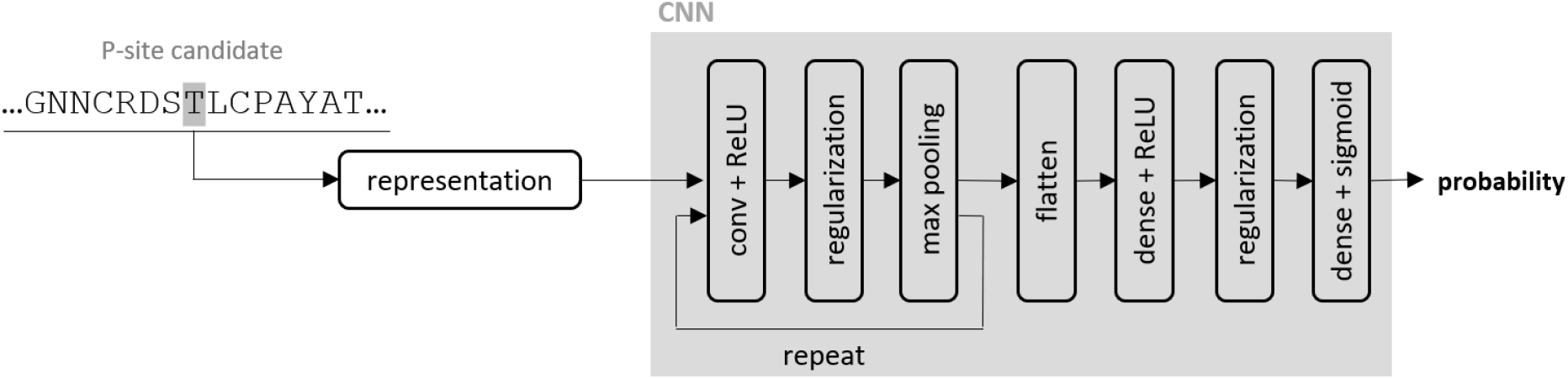
The CNN blueprint, highlighting the different components that are used throughout this study.

#### Benchmark methods

To evaluate the performance of the pLM-based predictors, we compare them to a one-hot encoded baseline, and to three state-of-the-art predictors: MusiteDeep^18^, DeepPhos^19^, and DeepPSP^13^. Code from their respective GitHub repositories was used to train their predictors from scratch on different data sources.

MusiteDeep implements two prediction modules: a CNN module enhanced with attention layers, and a capsule network module with dynamic routing, both built on a one-hot encoding with a receptive field of size 33 centered around the candidate phosphosite. Predicted probabilities for both modules are averaged to obtain the final phosphosite predictions.

In DeepPhos, three different densely connected CNN blocks are used with one-hot encoded inputs with receptive fields of size 15, 31 and 51, where each block consists of convolutional and intra-block concatenation layers. Blocks are combined using an inter-block concatenation layer, followed by a final fully connected layer.

DeepPSP uses two modules, each consisting of a convolutional block, a squeeze-and-excitation network, a bidirectional LSTM layer, and fully connected layers. The local module takes a one-hot encoding with a receptive field of length 51 in input, centered around the candidate phosphosite, while the global module uses a receptive field of length 2000 in input. Outputs of both modules are combined using a final fully connected layer.

Finally, LMPhosSite is the only approach that also uses pLMs in their pipeline. Like in DeepPSP, a local and a global module are considered. In the local module, a trainable embedding is used on a receptive field of 51 residues, followed by a CNN. The global module uses the ProtT5-XL-U50 embedding at the candidate phosphosite residue, followed by four fully-connected layers. Both models are trained individually, and the probabilities are then combined via stacked generalization, where a small neural network is trained to produce a final prediction. We reimplemented LMPhosSite in order to train the model on various datasets. In our implementation, we chose to use a learning rate of 1e-4 and batch size of 16, as these were not reported in the publication.

### Feature importance visualization

The fitted models are interpreted by applying SHapley Additive exPlanations^43^ (SHAP) to compute importance scores for individual residues in the receptive field that estimate their contribution to the predicted probability of a candidate phosphosite. For the visualization of all trained models we calculated the importance scores on the test set of the same dataset.

Importance scores are calculated using the DeepLiftSHAP implementation in the Captum^51^ software package. In DeepLiftSHAP, SHAP values are approximated relative to a reference baseline. For the baseline, we used the strategy that is proposed in the DeepLIFT paper^52^. Importance scores are calculated *n* times for each candidate phosphosite, where the baseline is obtained by shuffling the original sequence. Finally, the mean importance for each residue is calculated from those *n* runs. To keep the analysis computationally feasible, we used *n*=30. SHAP values are then normalized on a dataset level, such that the absolute values of all importance scores in one sequence add up to 100 on average.

The importance scores, also referred to as saliency maps, are computed on an individual base per candidate phosphosite. These maps are then aggregated into an average saliency map that we visualize as a sequence logo centered around the candidate phosphosite, cropped to contain ten positions both up- and downstream.

### Data acquisition

Prediction models were trained and evaluated on datasets that were compiled via three different setups. Firstly, we used the *multi-source* data that was employed in previous phosphosite prediction research, which we downloaded from the DeepPSP^13^ GitHub repository. This data was compiled from UniProtKB/SwissProt, dbPTM^40^, Phospho.ELM^41^, and PhosphoSitePlus^42^, with negative annotations defined as all remaining STY residues in the proteins with positive annotations. The multi-source data is also split up in separate ST and Y datasets. An important caveat with this standardized way of assembling phosphorylation datasets is that the negative set also contains STY residues for which phosphorylation can never be observed due to limitations in the data acquisition setups of MS/MS analysis.

Secondly, *Scop3P* phosphosite datasets were obtained from our in-house and publicly accessible Scop3P^7,39^ database, in which phosphosites were annotated from large-scale reprocessing of proteomics experiments^53^. All annotations were matched with protein sequences from UniProtKB/SwissProt (version 2021-11), discarding proteins when one of its annotations did not match with the sequence data. Non-phosphosite annotations were defined as all remaining STY residues in the proteins in the reprocessed experiments for which no evidence was found in PhosphoSitePlus or dbPTM.

Thirdly, *single-protease* datasets were obtained from individual experiments that involved different proteases^1^. In addition to individual datasets for each protease, we also compiled a combined dataset with annotations from the individual sets, which we labelled as *multi-protease*. Only ST annotations were considered, as Y annotations were too few. As before, negative labels were first defined as all ST residues in the annotated proteins that were not positively labelled in any of the single-protease experiments, Scop3P, or multi-source datasets.

Finally, datasets for other PTMs were obtained from PhosphositePlus and dbPTM. We considered the most abundant PTMs that were represented in both datasets, and combined annotations from both datasets. This led to further evaluation of our methodology on acetylation, methylation, sumoylation, and ubiquitination. A comprehensive overview of the datasets is given in Suppl. Tables 6 and 7.

### Peptide-filtering

As discussed before, negative annotations can then contain sites for which phosphorylation can never be observed with MS/MS experiments (e.g., sites in long stretches without peptide cleavage sites). To provide a higher quality dataset for both model training and evaluation, we propose an extra negative sample filtering step for datasets where experimental metadata is available. The filtering step rules out non-detectable sites from both training and evaluation by only including residues that occurred in matched peptide spectra (PSMs) in Scop3P. We will refer to this method as peptide filtering (PF) from here on. An overview of peptide-filtered datasets is given in Suppl. Table 8. Note that contrary to the non-peptide-filtered single- and multi-protease datasets, proteins without positive annotations can be present in this data, as they might have been matched with in the PSMs, hence increasing the number of proteins.

### Missed cleavage filtering

On top of peptide filtering, we considered an extra filtering step to further reduce the observed bias towards cleavage site residues. Instead of considering all peptides, we only retained those without missed cleavages. To be precise, we only keep PSMs for peptides that do not contain any R/K residues, unless they occur at the last position in the peptide. We refer to this method as PF-MC.

### Localization score filtering

Another attempted filtering step involved localization scores. For all candidate residues in the Scop3P-ST-PF dataset, we determined whether those residues were ever seen in a peptide with a non-localized phosphorylation across all phosphorylated peptides in the Scop3P metadata. As this would imply uncertainty of the “negative” annotation, we discarded residues for which this was the case from the negative set. We refer to this method as PF-Loc.

### Dataset selection in the different experiments

Datasets were selected and split for three main purposes in this manuscript: (a) model performance evaluation and comparison, (b) feature importance visualization, and (c) model transfer evaluation.

a. The comparison between one-hot encoded models, pLM-based models and existing predictors is done on the multi-source datasets, the Scop3P datasets, and the non-filtered multi-protease dataset. As PSMs in the single-protease datasets are obtained from individual experiments, their coverage is lower than in the multi-experiment Scop3P dataset, and thus, the number of non-phosphosite annotations is reduced drastically when applying peptide-filtering. This results in a smaller and much more balanced dataset (Suppl. Table 8), which impacts model training. Therefore, we did not consider these peptide-filtered datasets for comparison.
b. The visualization of important features is done by first training predictors on the datasets used for evaluation, and then calculating SHAP values on a fixed evaluation set. For models trained on ST data, we picked the test split of the multi-protease-PF (1,775 phosphosites, 2,331 non-phosphosites) for this purpose, given improved data quality via peptide filtering, while keeping computational costs low at the same time. For models trained on Y data, we calculated SHAP values using the Scop3P-Y-PF test set. For the other PTMs, SHAP values were computed on their respective test sets, where we limited each set to 7,000 candidate sites for the sake of computational feasibility. Note that we found that differences with experiments using different test sets for generating SHAP values were negligible.
c. The evaluation of model transferability was done on ST data only, to compare the Scop3P-ST-PF to the Scop3P-ST dataset, the multi-protease dataset, and the single-protease datasets. Evaluation was done on single- and multi-protease datasets.

### Dataset splits

For the multi-source data, the original data split between training and test was kept^13^, and training data was divided into an actual training part and a validation part. This is done for ST (Y) by randomly selecting 11,150 (7,999) proteins in the training set for training, and 738 (743) of all proteins for validation, while keeping the remaining 1,361 (968) proteins for testing. The Scop3P, single- and multi-protease datasets were randomly split by dividing proteins over training, validation and test sets via a 85/5/10 distribution. Finally, for the model transfer analysis, a cross-validation scheme was used, where each dataset set was split up into ten folds.

Then, for each fold, the proteins included in the validation set were removed from the training set, of which 1/10th was used for early stopping. These schemes are illustrated in Suppl. Fig. 10a and 10c. All datasets are made available via GitHub, along with their training/validation/test splits.

### Proteins are split into fragments

Due to computational complexity of transformer-based pLMs, all proteins were split into fragments of at most 512 amino acids before generating the representations. Fragments of one protein can be divided over multiple batches, but will always belong to the same training, validation, or test set to avoid data leakage.

Furthermore, as we wanted to avoid that a candidate phosphosite lies at the border between two fragments, thus implying that it can only use information from upstream or downstream residues, we allowed for overlapping fragments. Every 256 residues, a new fragment starts, and phosphosites are coupled to the fragment with the closest centre. The full batch split setup is illustrated in Suppl. Fig. 11.

For increased efficiency, we reduced the number of times the pLMs were used to generate representations. Per epoch, one protein fragment was forwarded through the pLM only once, even if it holds multiple phosphosite candidates. As a result, optimization was performed for a varying number of positive and negative phosphosites per batch.

### Hardware and software used

Programmatical frameworks used include PyTorch and PyTorch Lightning for model development and training and evaluation logic, WandB^54^ for experiment logging, and Captum^51^ for calculating the SHAP values. Sequence logos were created using LogoMaker^55^.

## Acknowledgements Funding

This work was supported by the Research Foundation – Flanders (FWO) [1274021N for J.Z.]; and the Vlaams Agentschap Innoveren en Ondernemen [HBC.2020.2205 for R.B.]. Research at our lab is core-funded by the VIB - Center for Medical Biotechnology and Ghent University.

