## Supplementary Materials for "A study on experimental bias in post-translational modification predictors"

|  | **Scop3P-ST** | | | **Scop3P-Y** | | |
| --- | --- | --- | --- | --- | --- | --- |
| **representation** | **precision for recall at 0.8** | **AUPRC** | **AUROC** | **precision for recall at 0.8** | **AUPRC** | **AUROC** |
| MusiteDeep | 0.396 (0.003) | 0.607 (0.003) | 0.935 (0.001) | 0.161 (0.003) | 0.301 (0.008) | 0.895 (0.001) |
| DeepPhos | 0.396 (0.011) | 0.618 (0.009) | 0.936 (0.002) | 0.121 (0.008) | 0.289 (0.015) | 0.873 (0.007) |
| DeepPSP | 0.459 (0.009) | 0.671 (0.007) | 0.947 (0.001) | 0.164 (0.006) | 0.304 (0.014) | 0.905 (0.004) |
| LMPhosSite | 0.457 (0.011) | 0.683 (0.006) | 0.945 (0.002) | 0.182 (0.012) | 0.470 (0.019) | 0.913 (0.004) |
| one-hot | 0.401 (0.010) | 0.629 (0.004) | 0.936 (0.001) | 0.144 (0.007) | 0.341 (0.012) | 0.893 (0.003) |
| ESM-1_small_ | 0.437 (0.017) | 0.679 (0.011) | 0.941 (0.002) | 0.180 (0.017) | 0.422 (0.038) | 0.908 (0.009) |
| ESM-1b | 0.427 (0.016) | 0.681 (0.008) | 0.936 (0.003) | 0.184 (0.016) | 0.464 (0.034) | 0.903 (0.006) |
| ESM-2_150M_ | 0.443 (0.018) | 0.685 (0.010) | 0.940 (0.004) | 0.184 (0.011) | 0.478 (0.022) | 0.911 (0.005) |
| ESM-2_650M_ | 0.439 (0.006) | 0.684 (0.004) | 0.939 (0.002) | 0.171 (0.007) | 0.474 (0.021) | 0.903 (0.003) |
| ESM-2_3B_ | 0.426 (0.019) | 0.679 (0.010) | 0.936 (0.003) | 0.162 (0.018) | 0.461 (0.019) | 0.900 (0.008) |
| CARP_640M_ | 0.393 (0.013) | 0.653 (0.010) | 0.932 (0.004) | 0.168 (0.013) | 0.441 (0.037) | 0.902 (0.008) |
| ProtT5_XL_U50 | **0.487 (0.014)** | **0.718 (0.005)** | **0.948 (0.003)** | **0.206 (0.007)** | **0.512 (0.018)** | **0.922 (0.006)** |
| Ankh_base | 0.449 (0.011) | 0.690 (0.009) | 0.941 (0.002) | 0.165 (0.015) | 0.470 (0.030) | 0.902 (0.005) |
| Ankh_large | 0.449 (0.008) | 0.690 (0.006) | 0.941 (0.002) | 0.200 (0.015) | 0.493 (0.018) | 0.917 (0.005) |
|  | **multi-source-ST** | | | **multi-source-Y** | | |
| **representation** | **precision for recall at 0.8** | **AUPRC** | **AUROC** | **precision for recall at 0.8** | **AUPRC** | **AUROC** |
| MusiteDeep (reported) | - | 0.46 | 0.80 | - | 0.33 | 0.69 |
| DeepPSP (reported) | - | 0.50 | 0.82 | - | 0.39 | 0.73 |
| LMPhosSite (reported) | - | 0.532 | **0.823** | - | **0.443** | **0.747** |
| MusiteDeep (retrained) | 0.270 (0.001) | 0.452 (0.001) | 0.789 (0.001) | 0.249 (0.002) | 0.387 (0.008) | 0.709 (0.002) |
| DeepPhos (retrained) | 0.282 (0.001) | 0.456 (0.005) | 0.800 (0.001) | 0.248 (0.007) | 0.348 (0.011) | 0.696 (0.009) |
| DeepPSP (retrained) | 0.289 (0.003) | 0.485 (0.007) | 0.810 (0.003) | 0.268 (0.003) | 0.389 (0.004) | 0.731 (0.003) |
| LMPhosSite (retrained) | **0.292 (0.002)** | 0.521 (0.003) | 0.812 (0.001) | 0.259 (0.002) | 0.442 (0.004) | 0.731 (0.003) |
| one-hot | 0.284 (0.005) | 0.475 (0.008) | 0.805 (0.005) | 0.255 (0.004) | 0.366 (0.010) | 0.711 (0.008) |
| ESM-1_small_ | 0.284 (0.008) | 0.520 (0.007) | 0.808 (0.007) | 0.262 (0.004) | 0.421 (0.010) | 0.728 (0.005) |
| ESM-1b | 0.270 (0.009) | 0.514 (0.009) | 0.796 (0.007) | 0.265 (0.005) | **0.443 (0.004)** | **0.737 (0.005)** |
| ESM-2_150M_ | 0.280 (0.008) | 0.514 (0.011) | 0.803 (0.007) | 0.255 (0.007) | 0.418 (0.012) | 0.723 (0.009) |
| ESM-2_650M_ | 0.276 (0.009) | 0.510 (0.012) | 0.800 (0.009) | 0.253 (0.008) | 0.427 (0.025) | 0.720 (0.015) |
| ESM-2_3B_ | 0.266 (0.009) | 0.501 (0.010) | 0.791 (0.009) | 0.247 (0.010) | 0.412 (0.020) | 0.711 (0.016) |
| CARP_640M_ | 0.279 (0.004) | 0.510 (0.006) | 0.802 (0.003) | 0.261 (0.003) | 0.436 (0.006) | 0.731 (0.004) |
| ProtT5_XL_U50 | **0.292 (0.008)** | **0.544 (0.007)** | **0.817 (0.006)** | 0.262 (0.005) | 0.442 (0.016) | 0.734 (0.010) |
| Ankh_base | 0.275 (0.005) | 0.512 (0.007) | 0.801 (0.005) | 0.258 (0.008) | 0.426 (0.021) | 0.726 (0.012) |
| Ankh_large | 0.280 (0.005) | 0.515 (0.007) | 0.804 (0.003) | 0.257 (0.010) | 0.426 (0.021) | 0.724 (0.015) |
|  | **multi-protease** | | |  |  |  |
| **representation** | **precision for recall at 0.8** | **AUPRC** | **AUROC** |  |  |  |
| MusiteDeep | 0.221 (0.003) | 0.498 (0.006) | 0.913 (0.001) |  |  |  |
| DeepPhos | 0.191 (0.009) | 0.498 (0.013) | 0.904 (0.003) |  |  |  |
| DeepPSP | 0.236 (0.016) | 0.543 (0.007) | 0.919 (0.003) |  |  |  |
| LMPhosSite | 0.232 (0.011) | 0.585 (0.009) | 0.915 (0.002) |  |  |  |
| one-hot | 0.189 (0.012) | 0.492 (0.009) | 0.903 (0.004) |  |  |  |
| ESM-1_small_ | 0.254 (0.023) | 0.592 (0.010) | 0.917 (0.008) |  |  |  |
| ESM-1b | 0.214 (0.017) | 0.575 (0.012) | 0.905 (0.007) |  |  |  |
| ESM-2_150M_ | 0.240 (0.011) | 0.588 (0.008) | 0.916 (0.005) |  |  |  |
| ESM-2_650M_ | 0.236 (0.020) | 0.591 (0.009) | 0.912 (0.006) |  |  |  |
| ESM-2_3B_ | 0.194 (0.011) | 0.556 (0.011) | 0.901 (0.004) |  |  |  |
| CARP_640M_ | 0.207 (0.023) | 0.569 (0.009) | 0.908 (0.008) |  |  |  |
| ProtT5_XL_U50 | **0.241 (0.017)** | **0.603 (0.010)** | **0.919 (0.005)** |  |  |  |
| Ankh_base | 0.215 (0.012) | 0.579 (0.005) | 0.908 (0.005) |  |  |  |
| Ankh_large | 0.234 (0.013) | 0.584 (0.008) | 0.914 (0.003) |  |  |  |

**Supplementary Table 1.** Model performance for phosphorylation prediction on the Scop3P, multi-source, and multi-protease datasets. Average metrics and standard deviations over 10 separate runs are reported.

|  | **Acetylation** | | | **Methylation** | | |
| --- | --- | --- | --- | --- | --- | --- |
| **representation** | **precision for recall at 0.8** | **AUPRC** | **AUROC** | **precision for recall at 0.8** | **AUPRC** | **AUROC** |
| one-hot | 0.148 (0.005) | 0.266 (0.021) | 0.754 (0.011) | 0.054 (0.005) | 0.171 (0.014) | 0.806 (0.016) |
| ProtT5-XL-U50 | 0.169 (0.002) | 0.365 (0.010) | 0.795 (0.004) | 0.062 (0.003) | 0.200 (0.019) | 0.831 (0.008) |
|  | **Sumoylation** | | | **Ubiquitination** | | |
| **representation** | **precision for recall at 0.8** | **AUPRC** | **AUROC** | **precision for recall at 0.8** | **AUPRC** | **AUROC** |
| one-hot | 0.151 (0.009) | 0.385 (0.041) | 0.796 (0.017) | 0.282 (0.007) | 0.397 (0.016) | 0.778 (0.008) |
| ProtT5-XL-U50 | 0.191 (0.010) | 0.480 (0.012) | 0.844 (0.006) | 0.300 (0.004) | 0.452 (0.004) | 0.801 (0.003) |

**Supplementary Table 2.** Model performance for prediction of other PTMs. Average metrics and standard deviations over 10 separate runs are reported.

|  | **Scop3P-ST-PF** | | | **Scop3P-Y-PF** | | |
| --- | --- | --- | --- | --- | --- | --- |
| **representation** | **precision for recall at 0.8** | **AUPRC** | **AUROC** | **precision for recall at 0.8** | **AUPRC** | **AUROC** |
| MusiteDeep | 0.495 (0.002) | 0.688 (0.002) | 0.911 (0.000) | 0.198 (0.006) | 0.383 (0.008) | 0.847 (0.003) |
| DeepPhos | 0.482 (0.007) | 0.677 (0.011) | 0.907 (0.002) | 0.163 (0.010) | 0.364 (0.023) | 0.824 (0.009) |
| DeepPSP | 0.531 (0.004) | 0.724 (0.009) | 0.921 (0.001) | 0.175 (0.009) | 0.336 (0.018) | 0.830 (0.008) |
| LMPhosSite | 0.567 (0.005) | 0.756 (0.003) | 0.927 (0.001) | 0.249 (0.007) | 0.542 (0.006) | 0.884 (0.003) |
| one-hot | 0.490 (0.007) | 0.686 (0.005) | 0.909 (0.002) | 0.187 (0.008) | 0.399 (0.008) | 0.847 (0.004) |
| ESM-1_small_ | 0.553 (0.011) | 0.748 (0.005) | 0.920 (0.003) | 0.228 (0.016) | 0.491 (0.038) | 0.868 (0.005) |
| ESM-1b | 0.558 (0.011) | 0.750 (0.011) | 0.921 (0.003) | 0.246 (0.010) | 0.526 (0.022) | 0.883 (0.004) |
| ESM-2_150M_ | 0.556 (0.010) | 0.753 (0.006) | 0.923 (0.003) | 0.230 (0.014) | 0.524 (0.009) | 0.874 (0.006) |
| ESM-2_650M_ | 0.550 (0.007) | 0.747 (0.008) | 0.921 (0.002) | 0.248 (0.024) | 0.531 (0.023) | 0.880 (0.007) |
| ESM-2_3B_ | 0.530 (0.006) | 0.743 (0.005) | 0.918 (0.001) | 0.230 (0.012) | 0.516 (0.018) | 0.872 (0.008) |
| CARP_640M_ | 0.518 (0.010) | 0.729 (0.007) | 0.915 (0.003) | 0.242 (0.012) | 0.521 (0.011) | 0.880 (0.004) |
| ProtT5_XL_U50 | **0.589 (0.010)** | **0.771 (0.008)** | **0.931 (0.002)** | **0.267 (0.013)** | **0.568 (0.014)** | **0.894 (0.005)** |
| Ankh_base | 0.555 (0.010) | 0.755 (0.008) | 0.923 (0.002) | 0.243 (0.015) | 0.550 (0.014) | 0.881 (0.006) |
| Ankh_large | 0.552 (0.007) | 0.743 (0.011) | 0.924 (0.002) | 0.252 (0.019) | 0.552 (0.017) | 0.888 (0.005) |

**Supplementary Table 3.** Model performance for phosphorylation prediction on the peptide-filtered Scop3P datasets. Average metrics and standard deviations over 10 separate runs are reported.

| (a) MAPK | (b) CDK | (c) PKa | (d) CK2 |
| --- | --- | --- | --- |
| 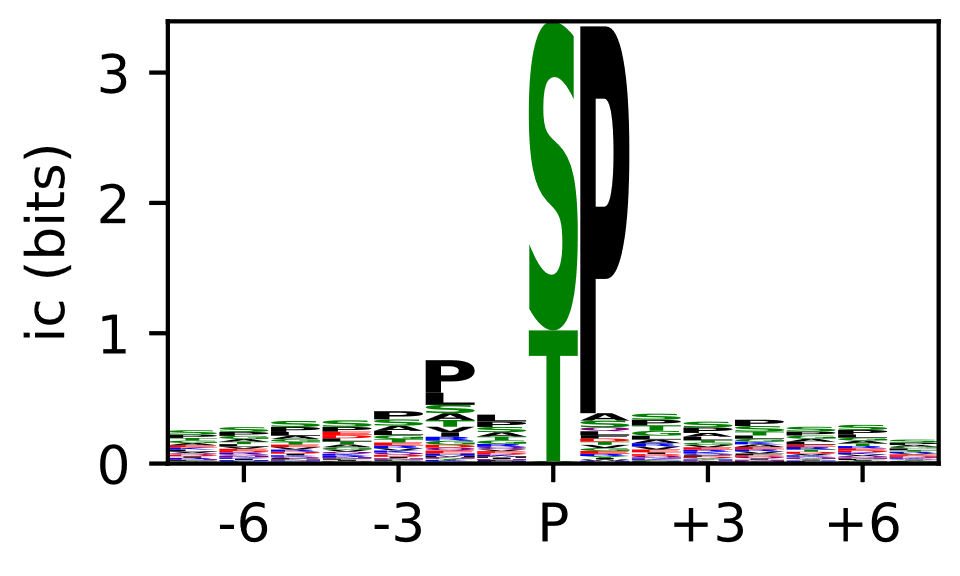 | 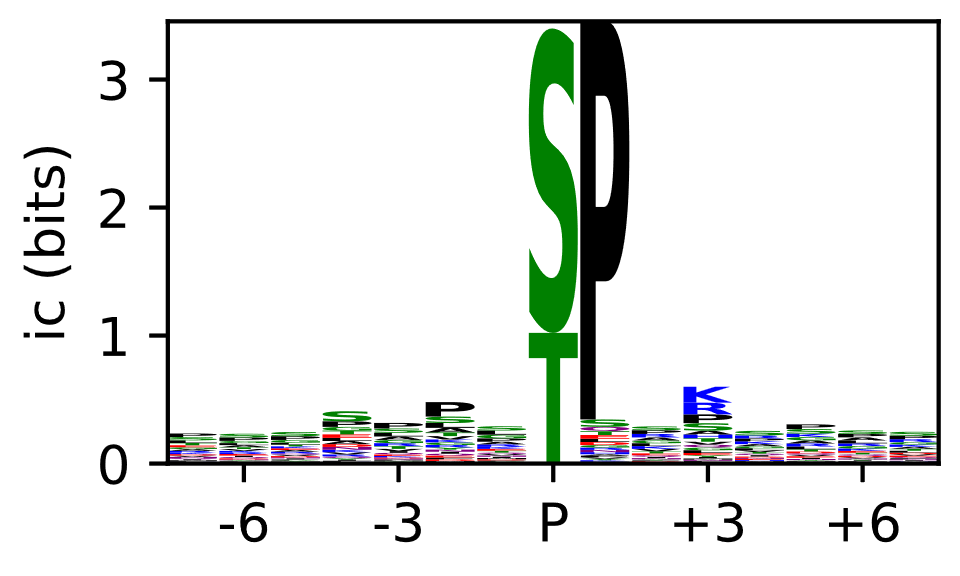 | 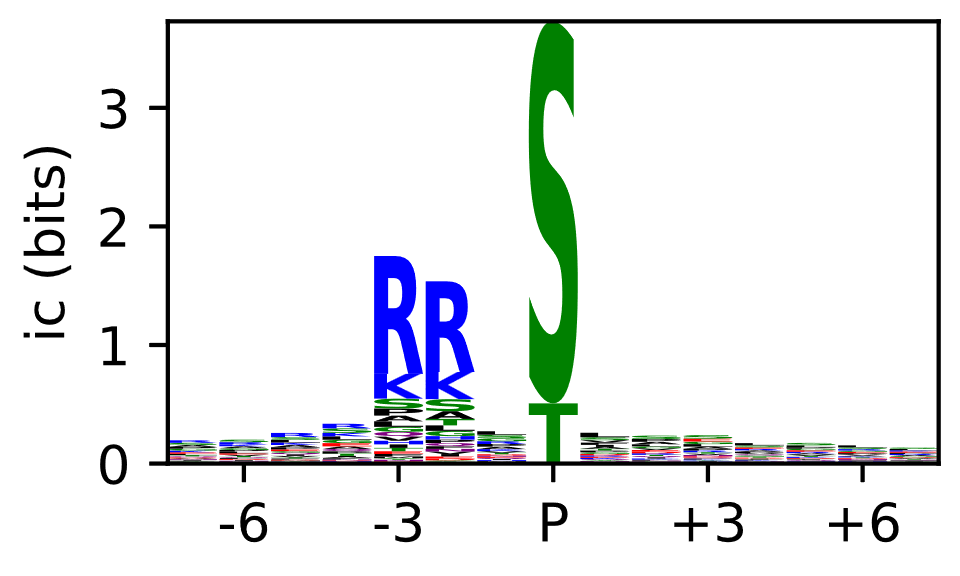 | 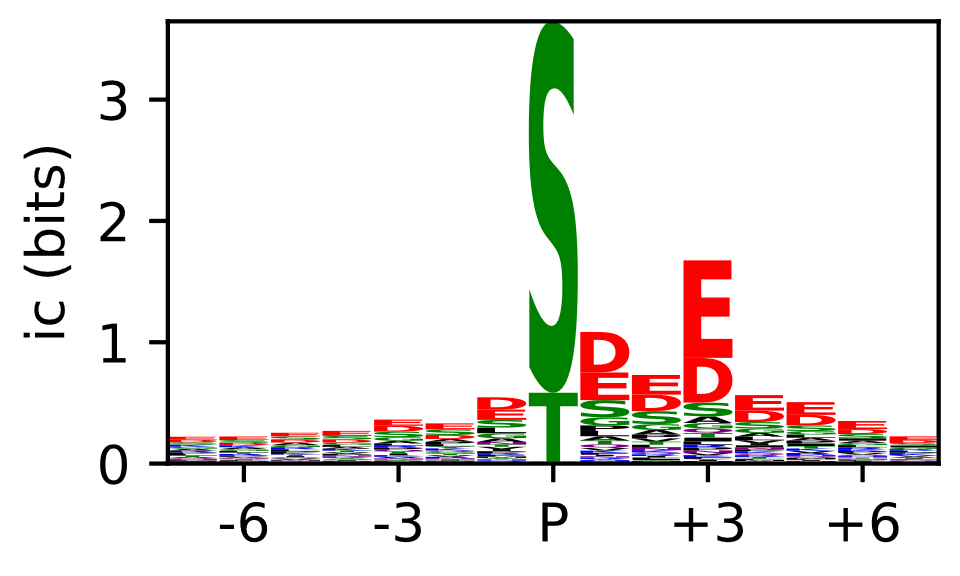 |

**Supplementary Figure 1.** Sequence logos, displaying information content (ic) per position, as derived from phosphosites with kinase annotations in PhosphoSitePlus^1^.

| one-hot, trained **on Scop3P-ST** data | one-hot, trained on **Scop3P-Y** data |
| --- | --- |
| 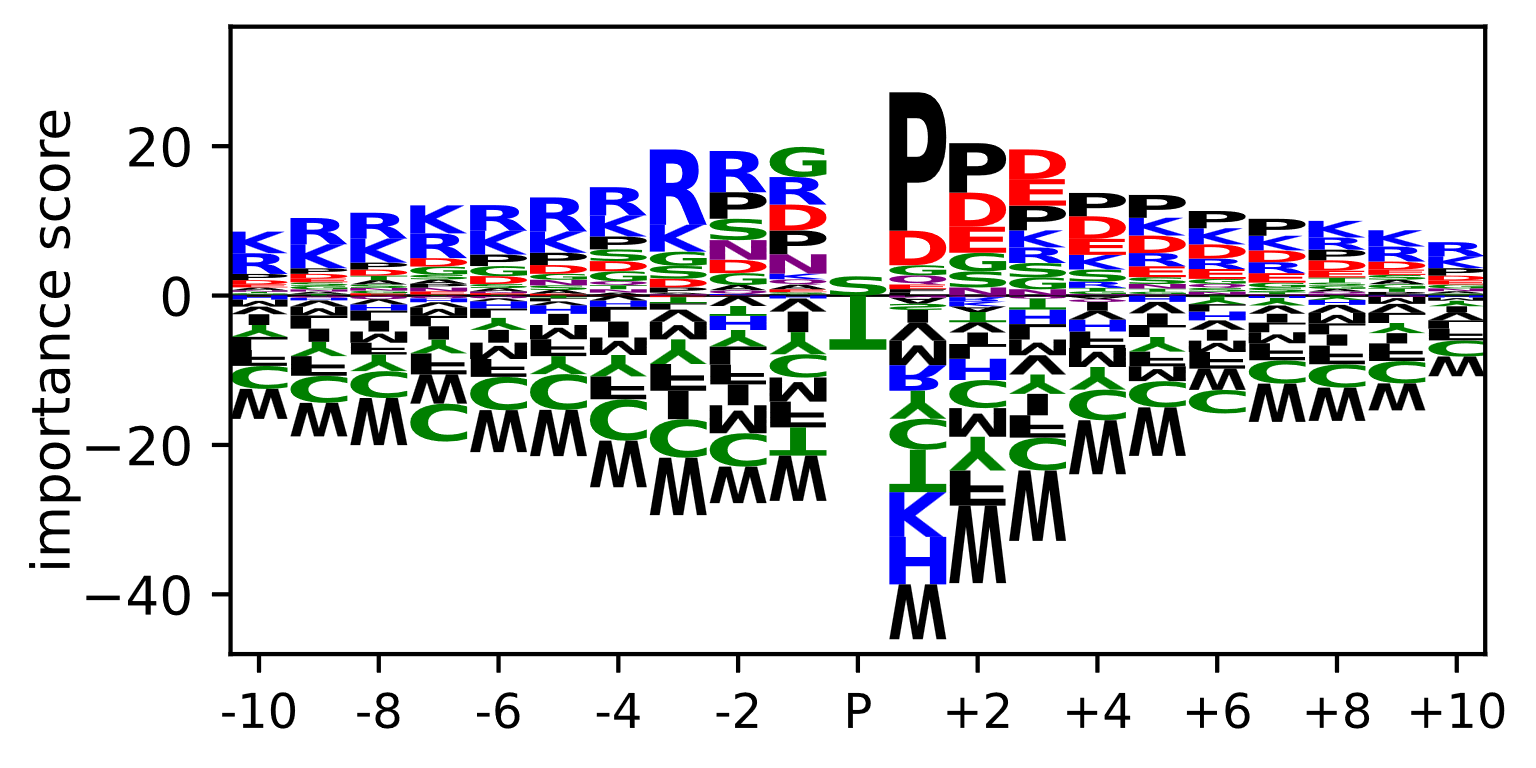 | 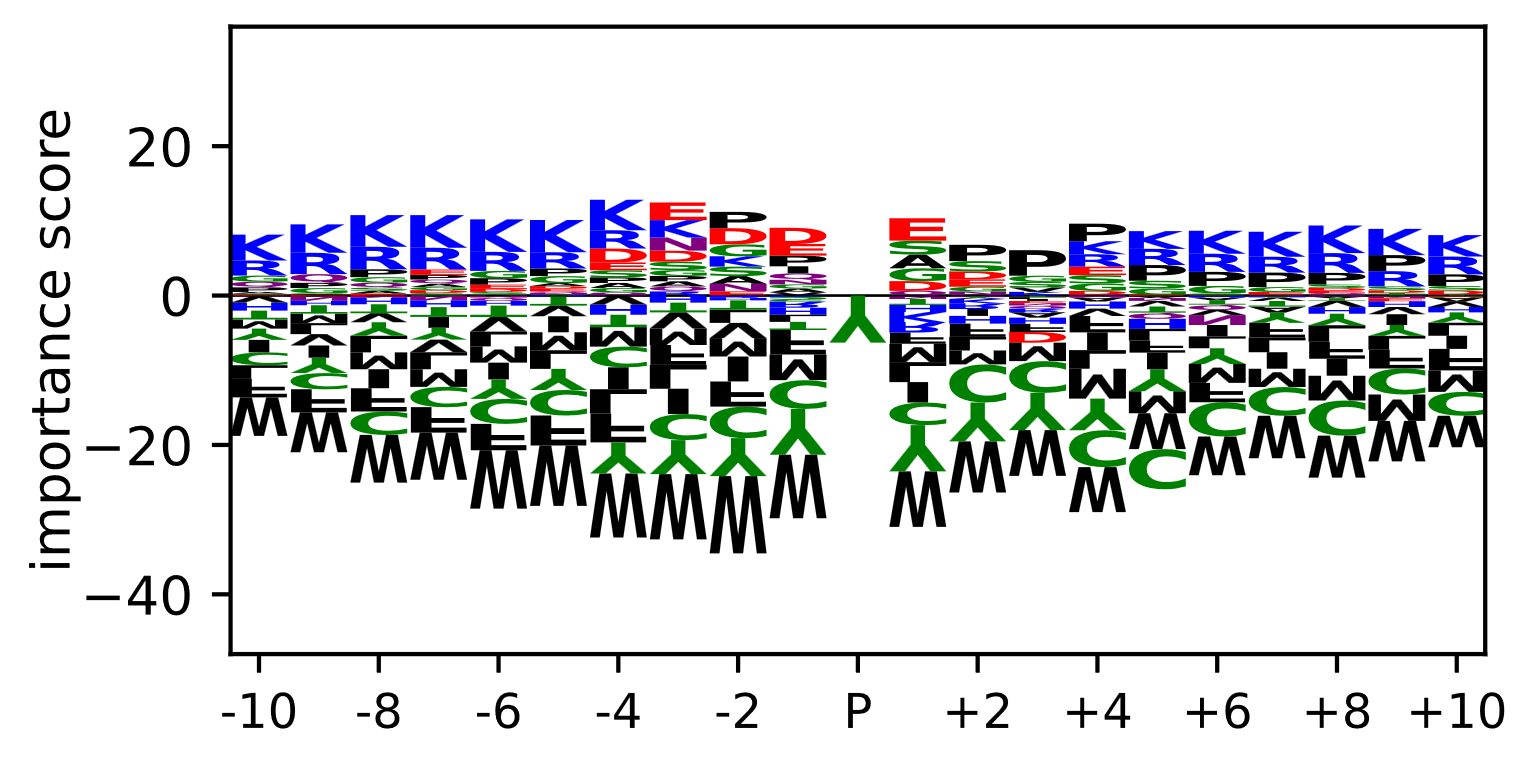 |

**Supplementary Figure 2.** Average importance scores per position, calculated using DeepLiftSHAP, for the one-hot encoded models on the Scop3P datasets.

| one-hot, trained on **Acetylation** data | one-hot, trained on **Methylation** data |
| --- | --- |
| 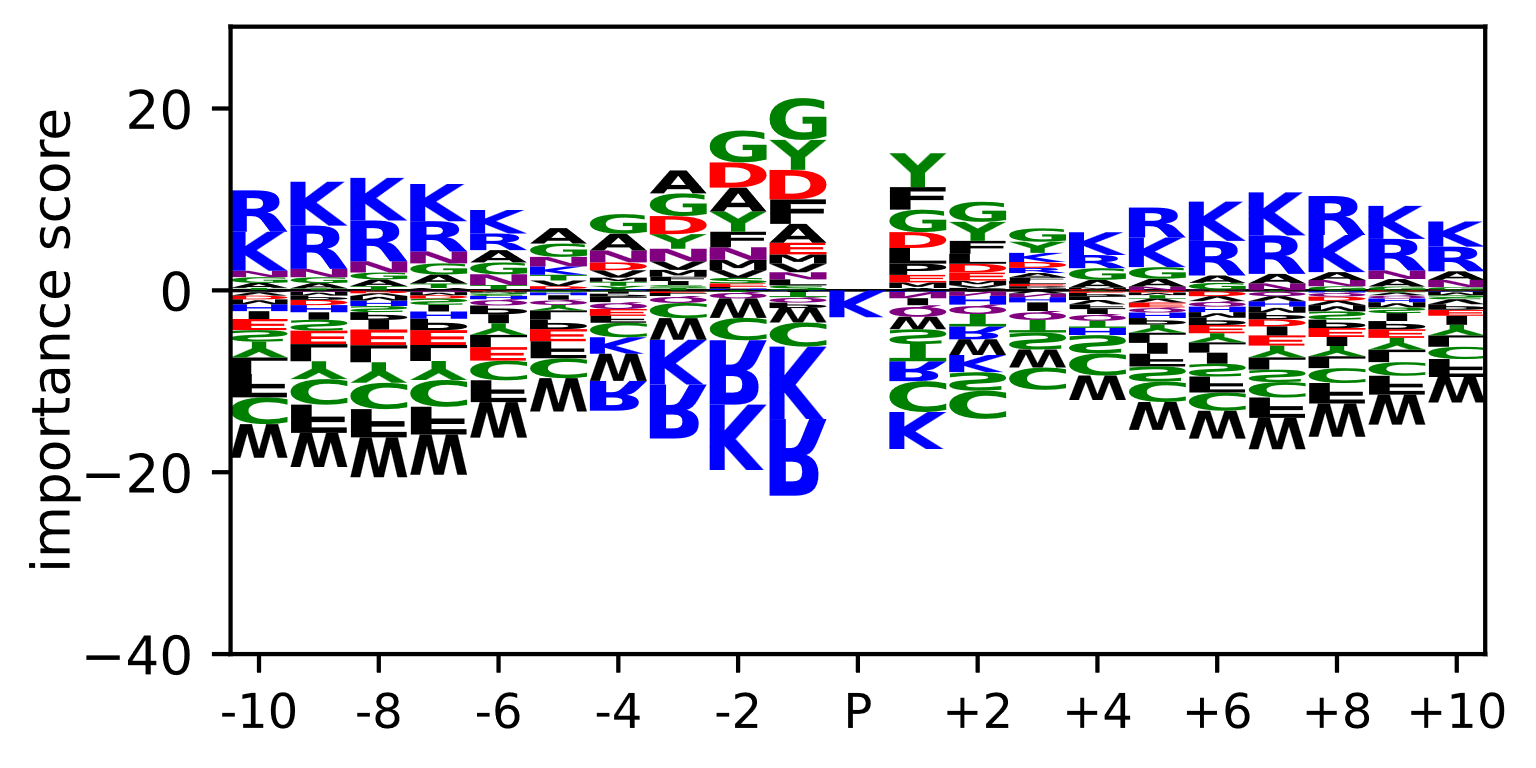 | 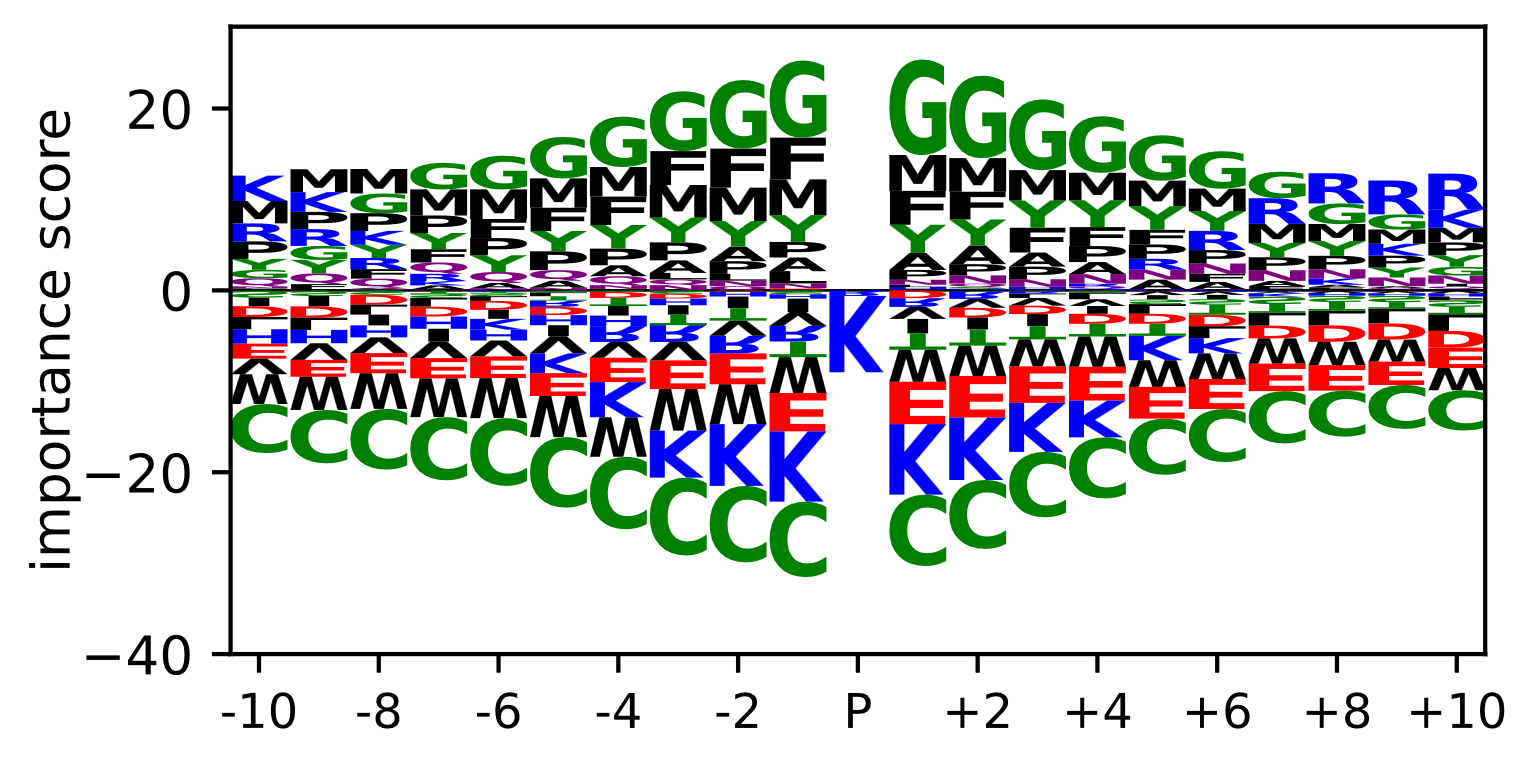 |
| one-hot, trained on **Sumoylation** data | one-hot, trained on **Ubiquitination** data |
| 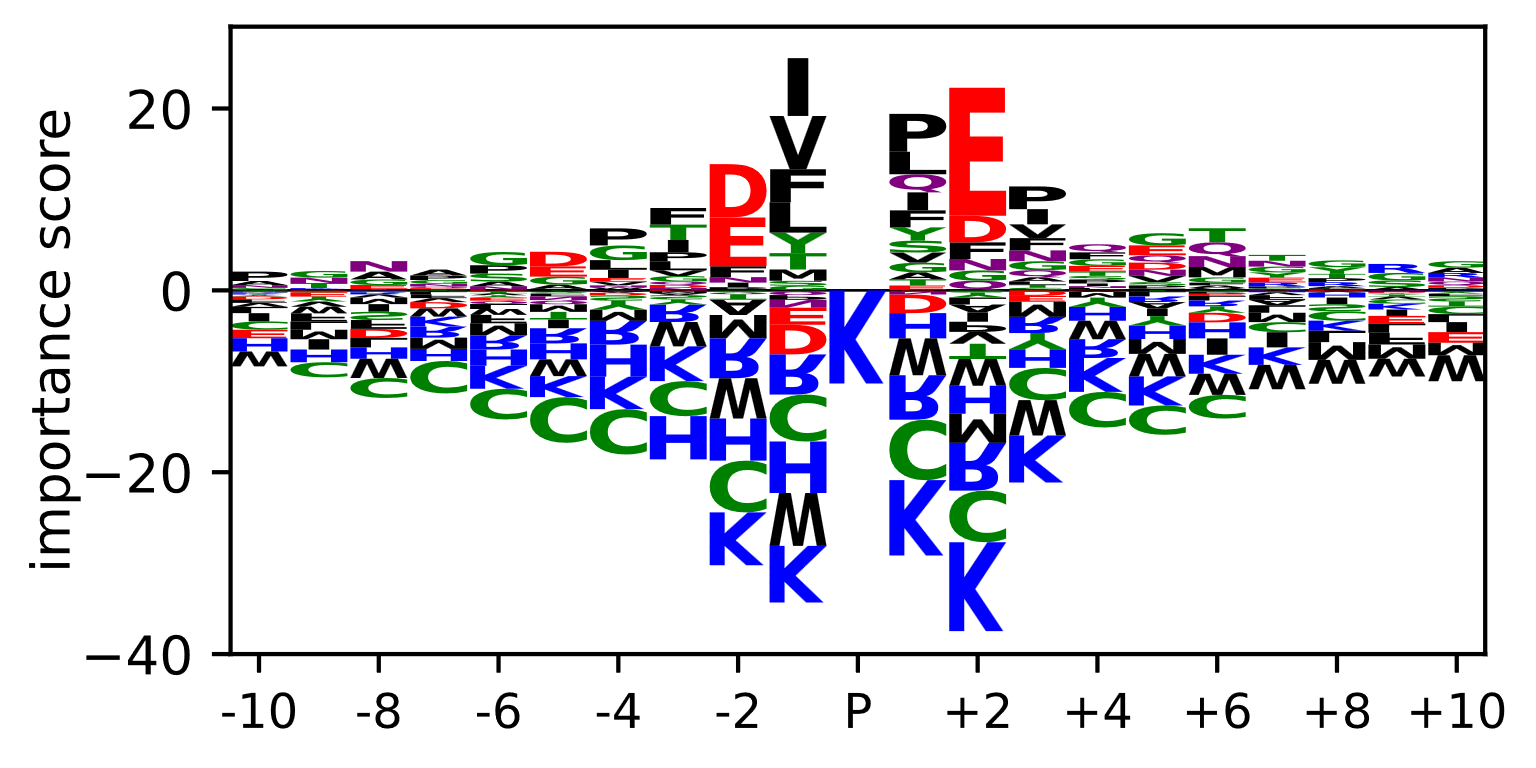 | 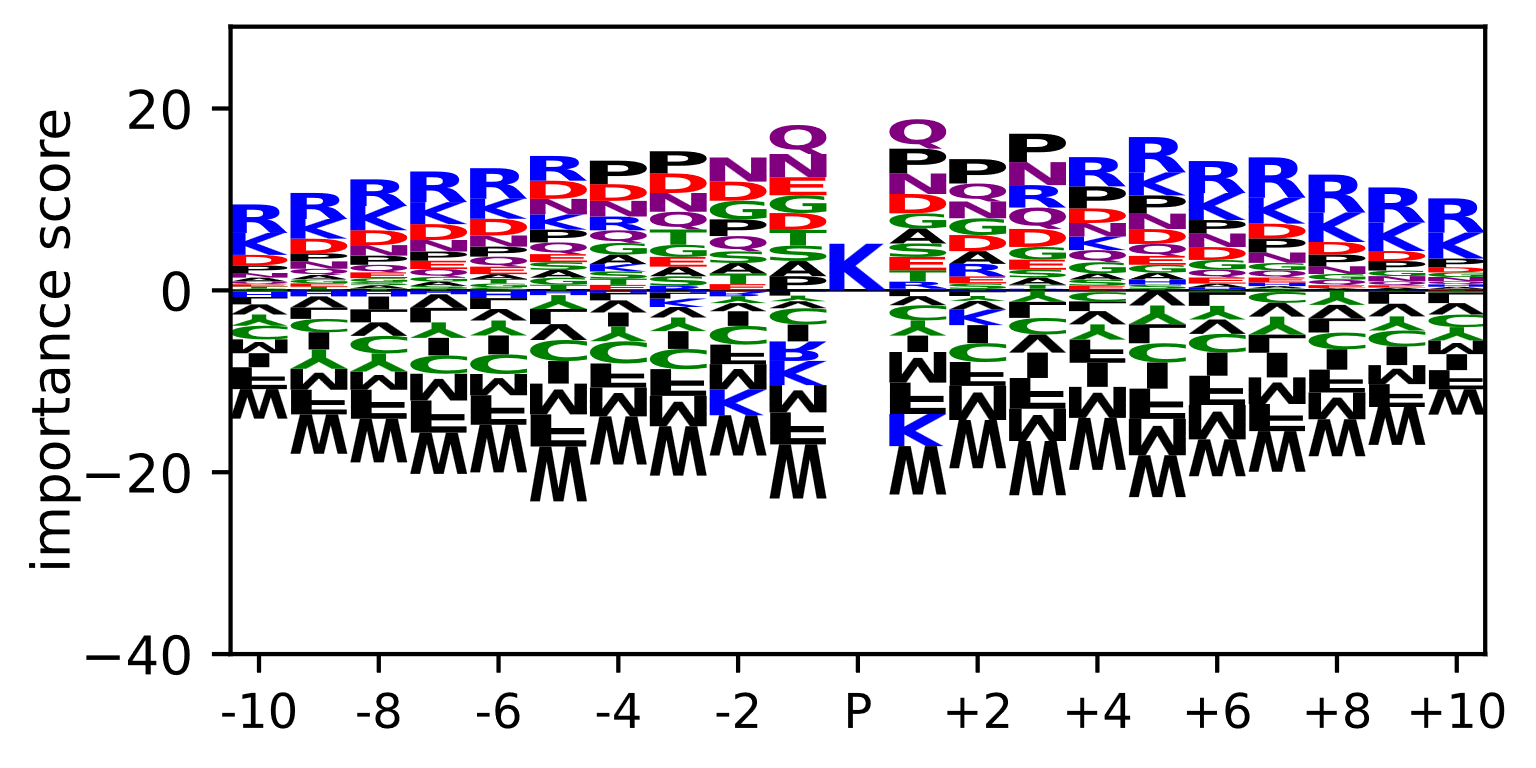 |

***Supplementary Figure 3.*** *Average importance scores per position, calculated using DeepLiftSHAP, for one-hot encoded an models trained on the acetylation, methylation, sumoylation, and ubiquitination datasets. Scores are calculated on their respective test sets.*

| one-hot, trained on **single-protease AspN** data | one-hot, trained on **single-protease chymotrypsin** data |
| --- | --- |
| 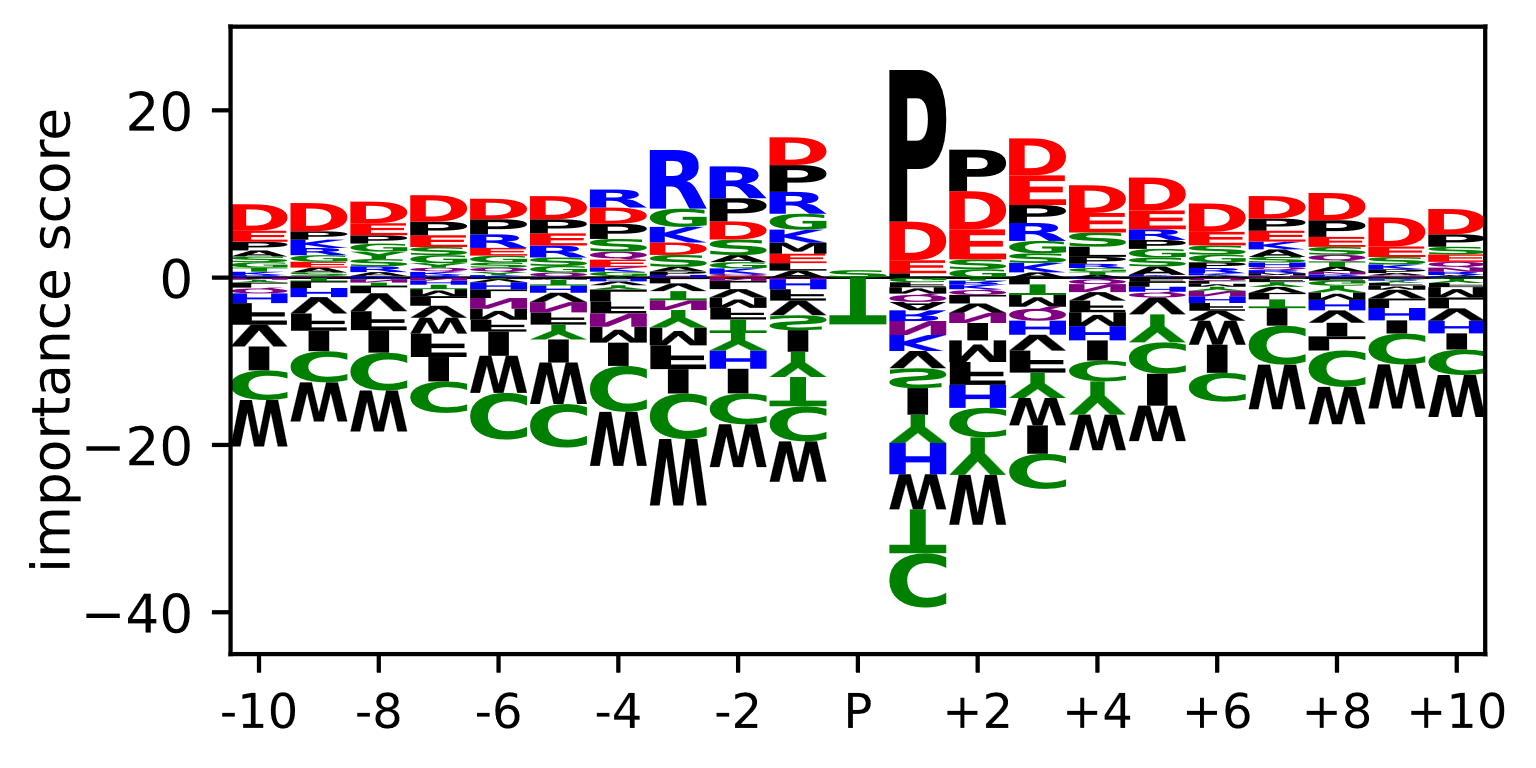 | 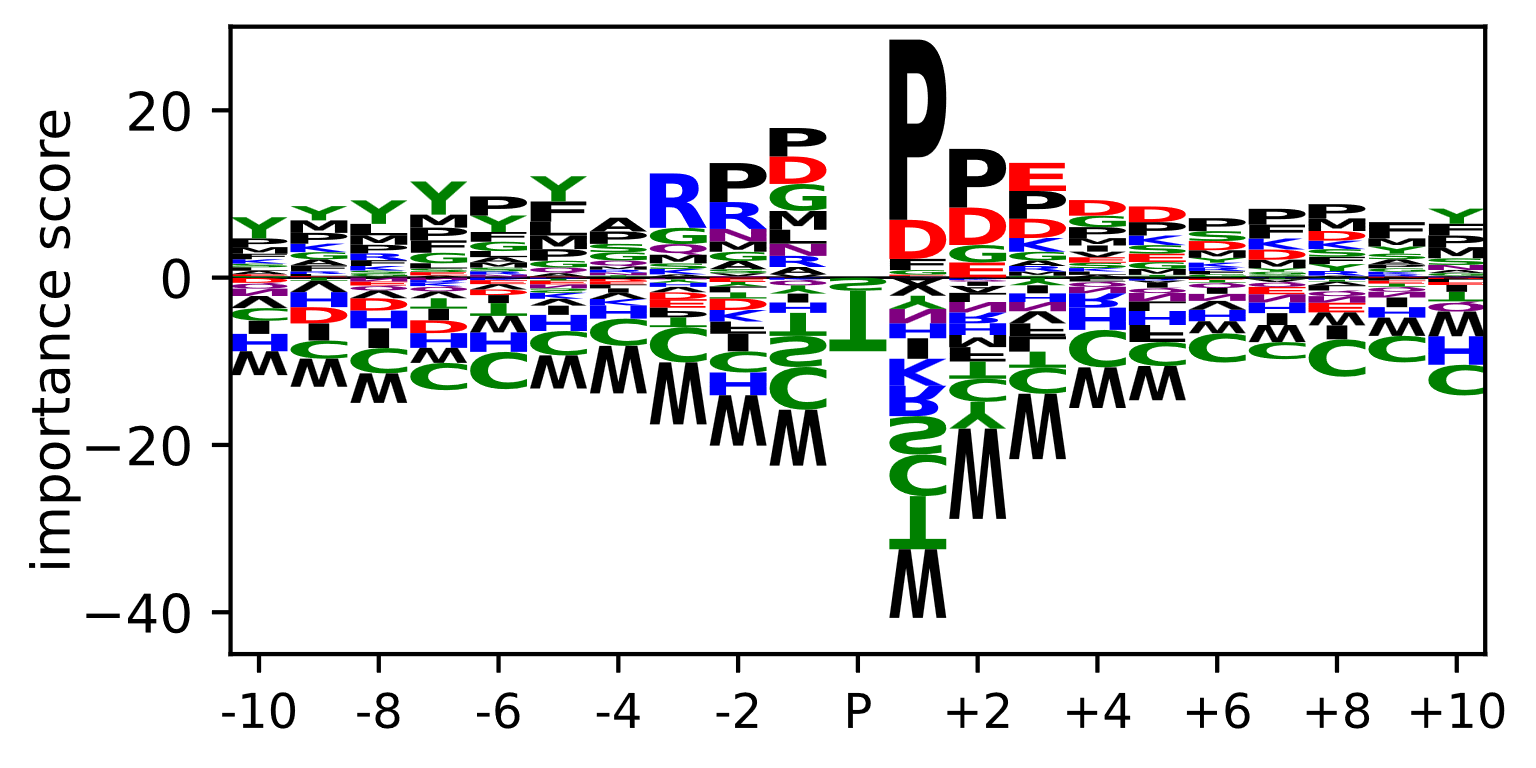 |
| one-hot, trained on **single-protease GluC** data | one-hot, trained on **single-protease LysC** data |
| 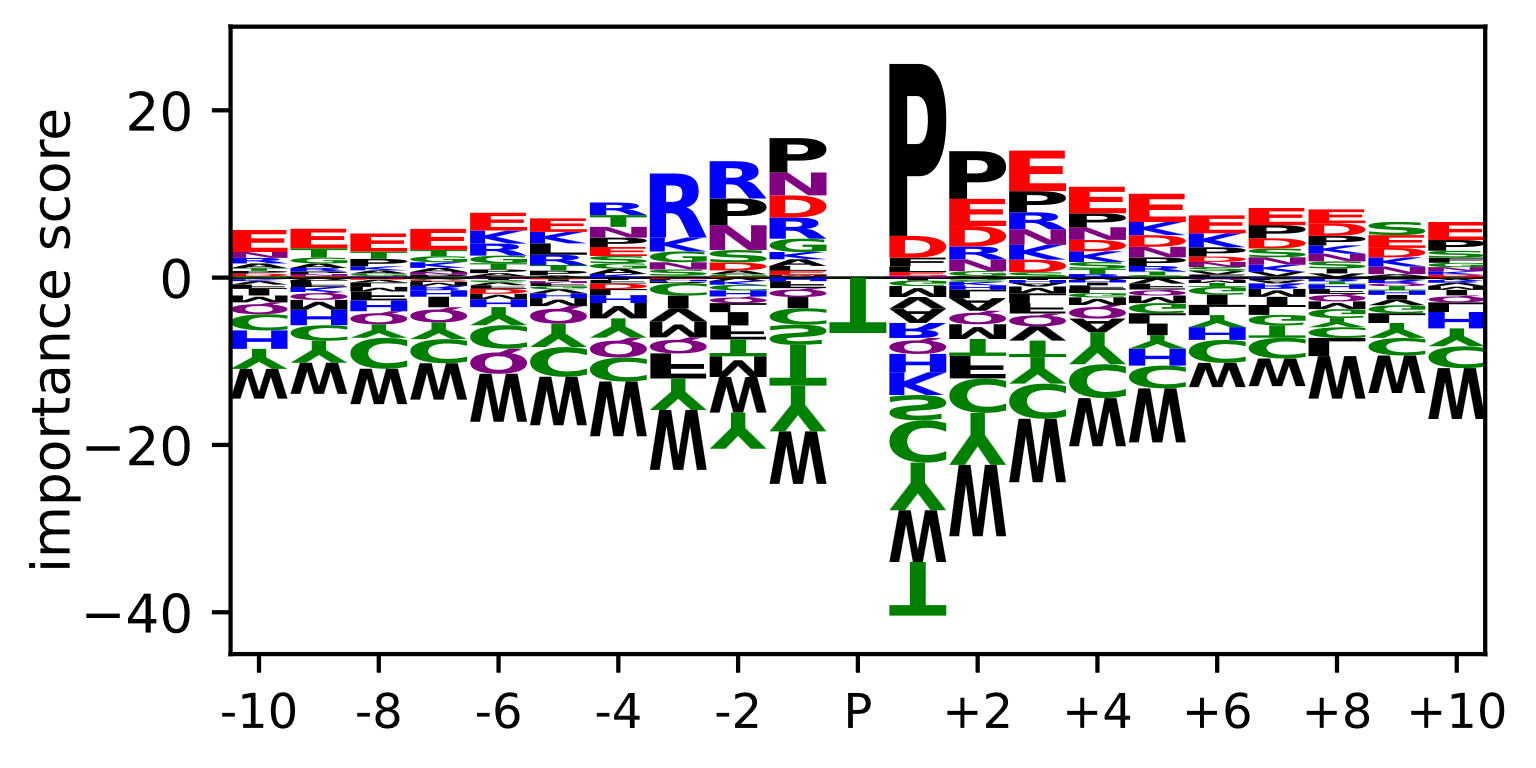 | 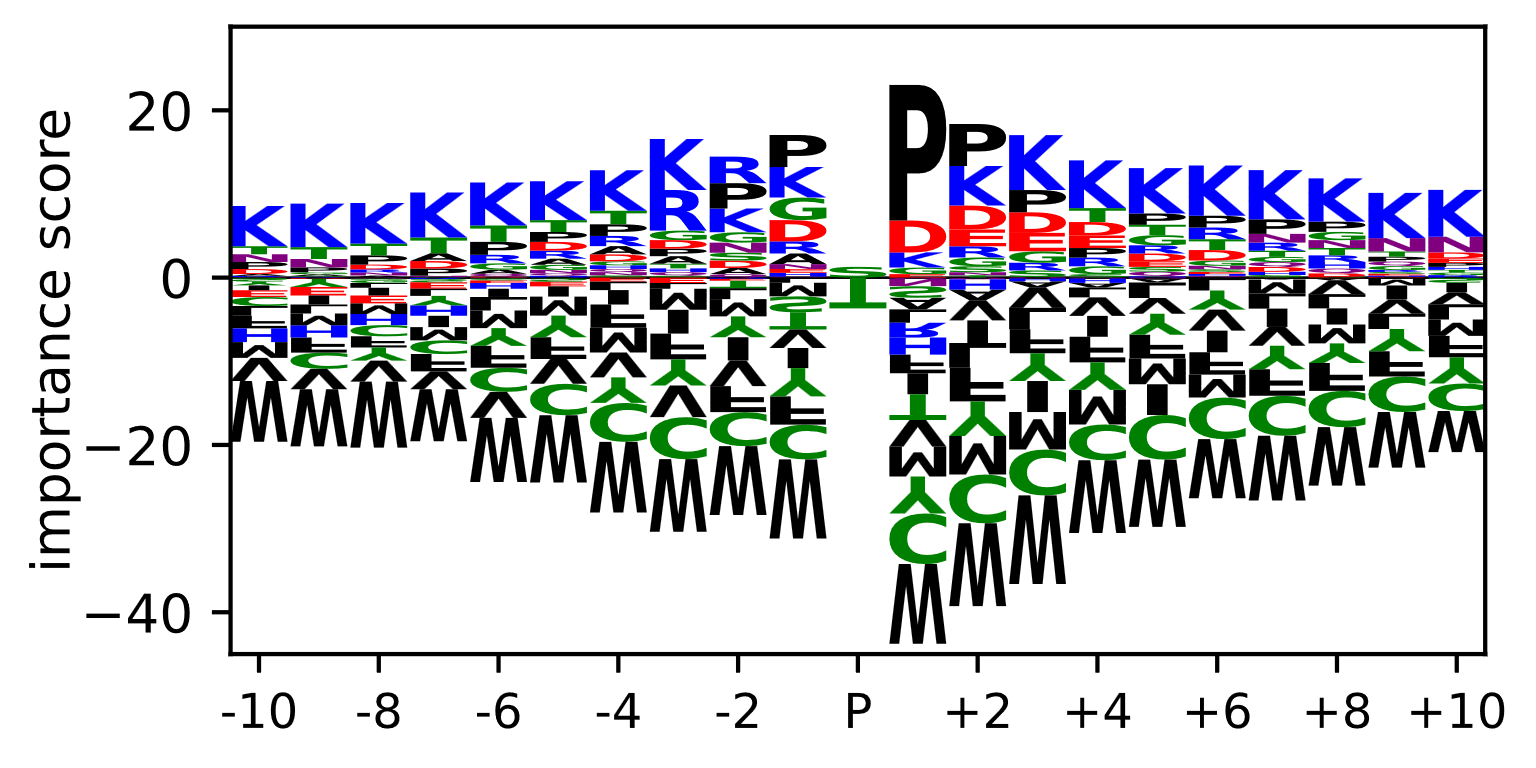 |
| one-hot, trained on **single-protease trypsin** data | one-hot, trained on **multi-protease** data |
| 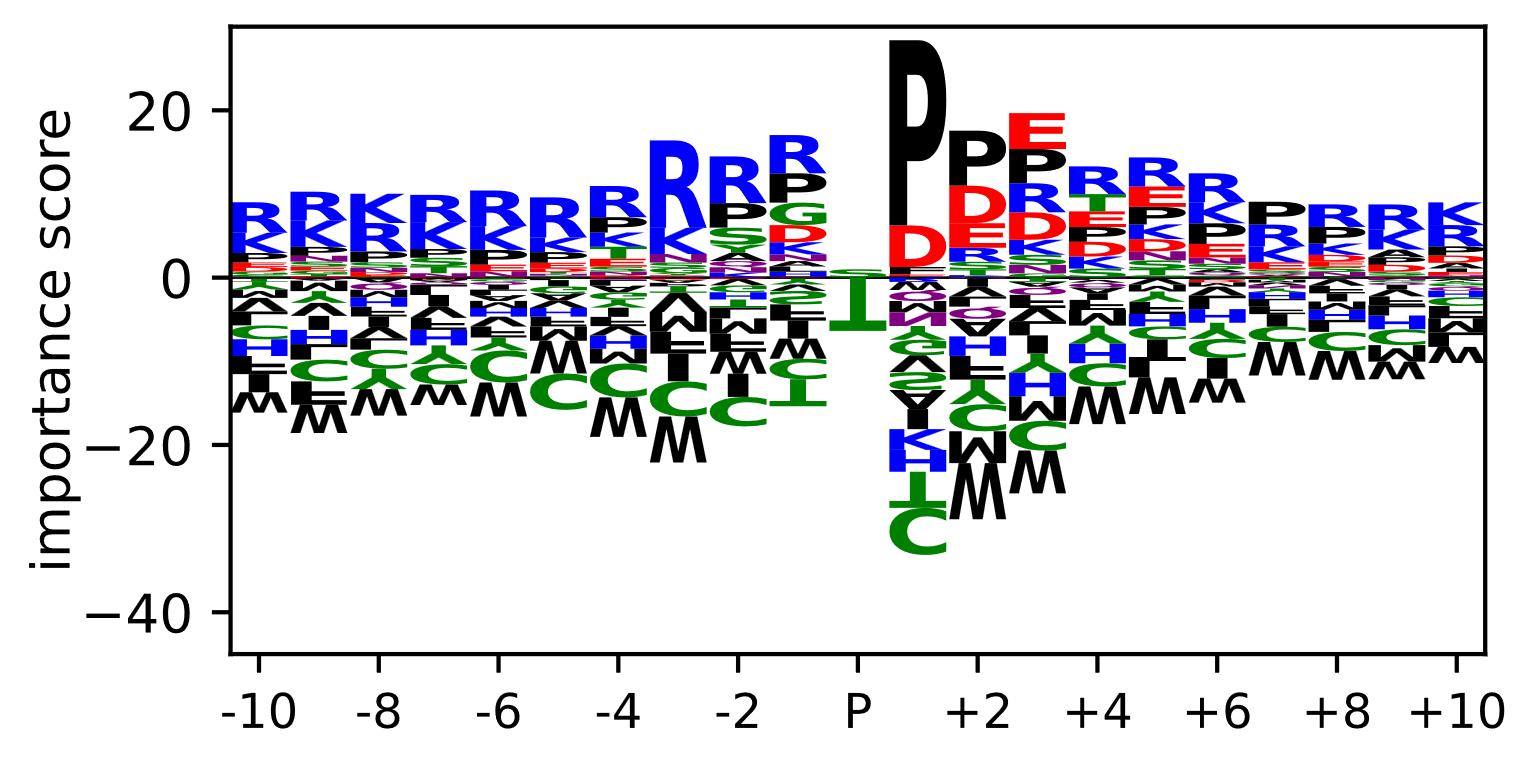 | 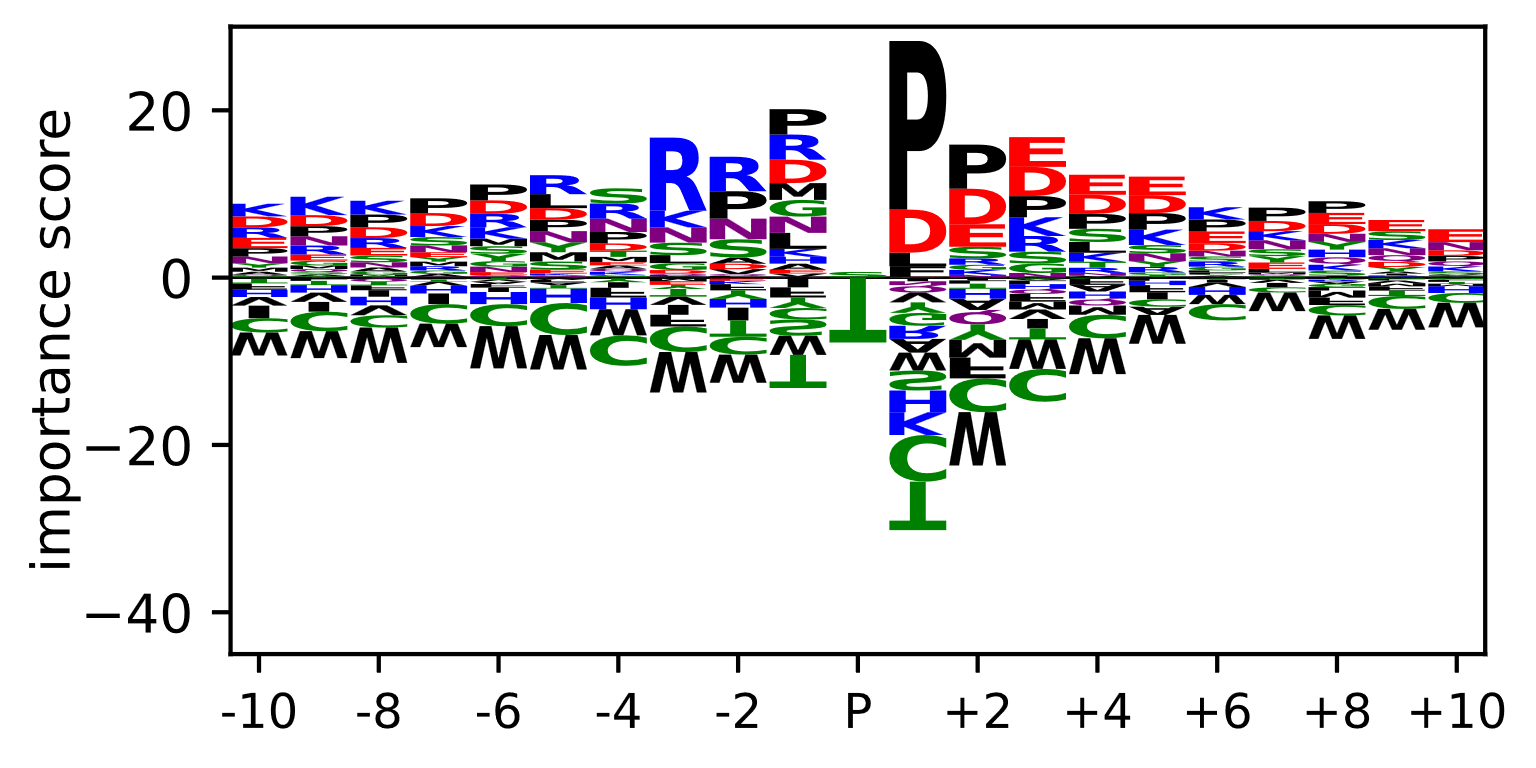 |

**Supplementary Figure 4.** Average importance scores per position, calculated using DeepLiftSHAP, for one-hot encoded models trained on the single-protease datasets, and the combined multi-protease dataset.

| one-hot, trained on **Scop3P-ST-PF** data | one-hot, trained on **Scop3P-Y-PF** data |
| --- | --- |
| 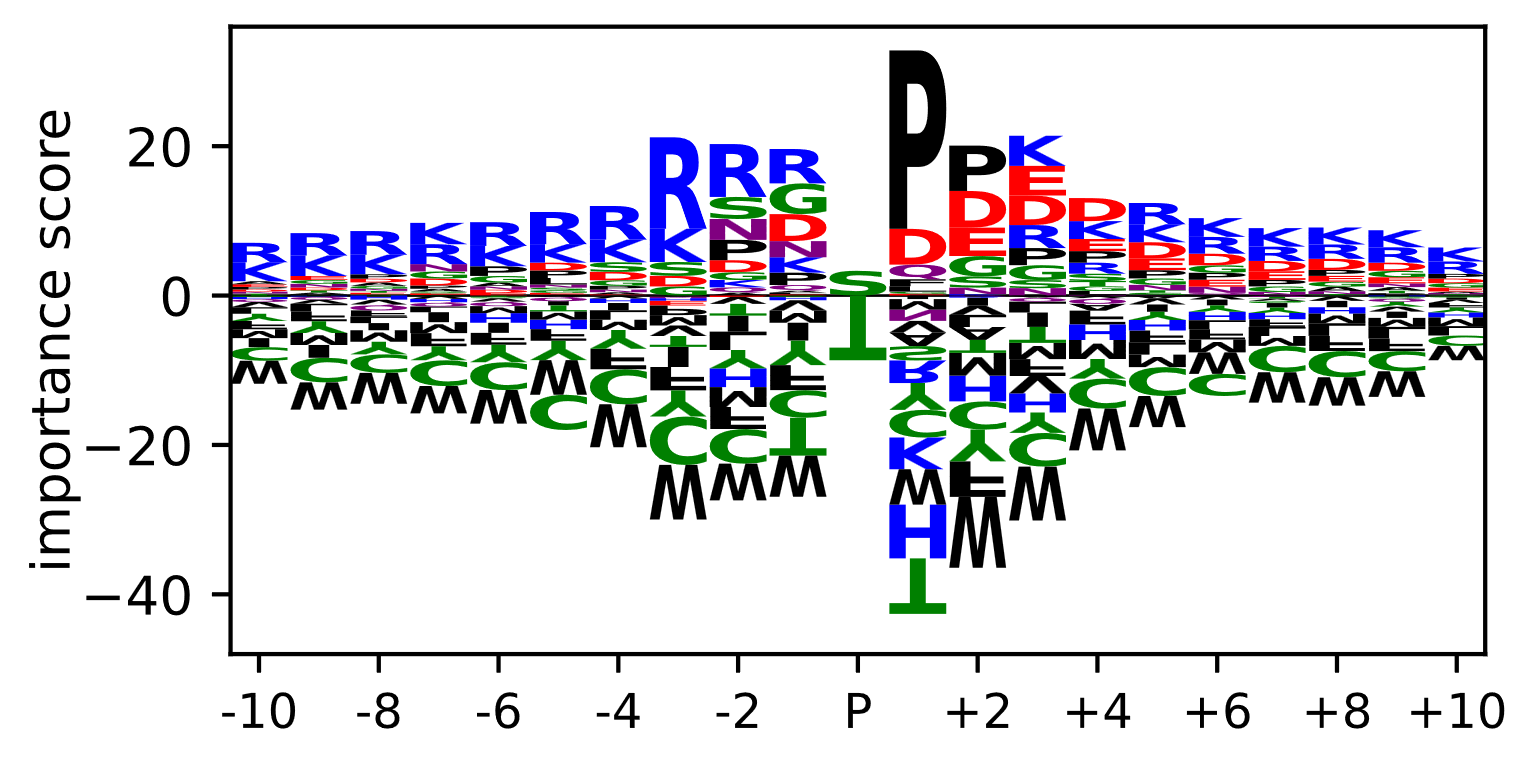 | 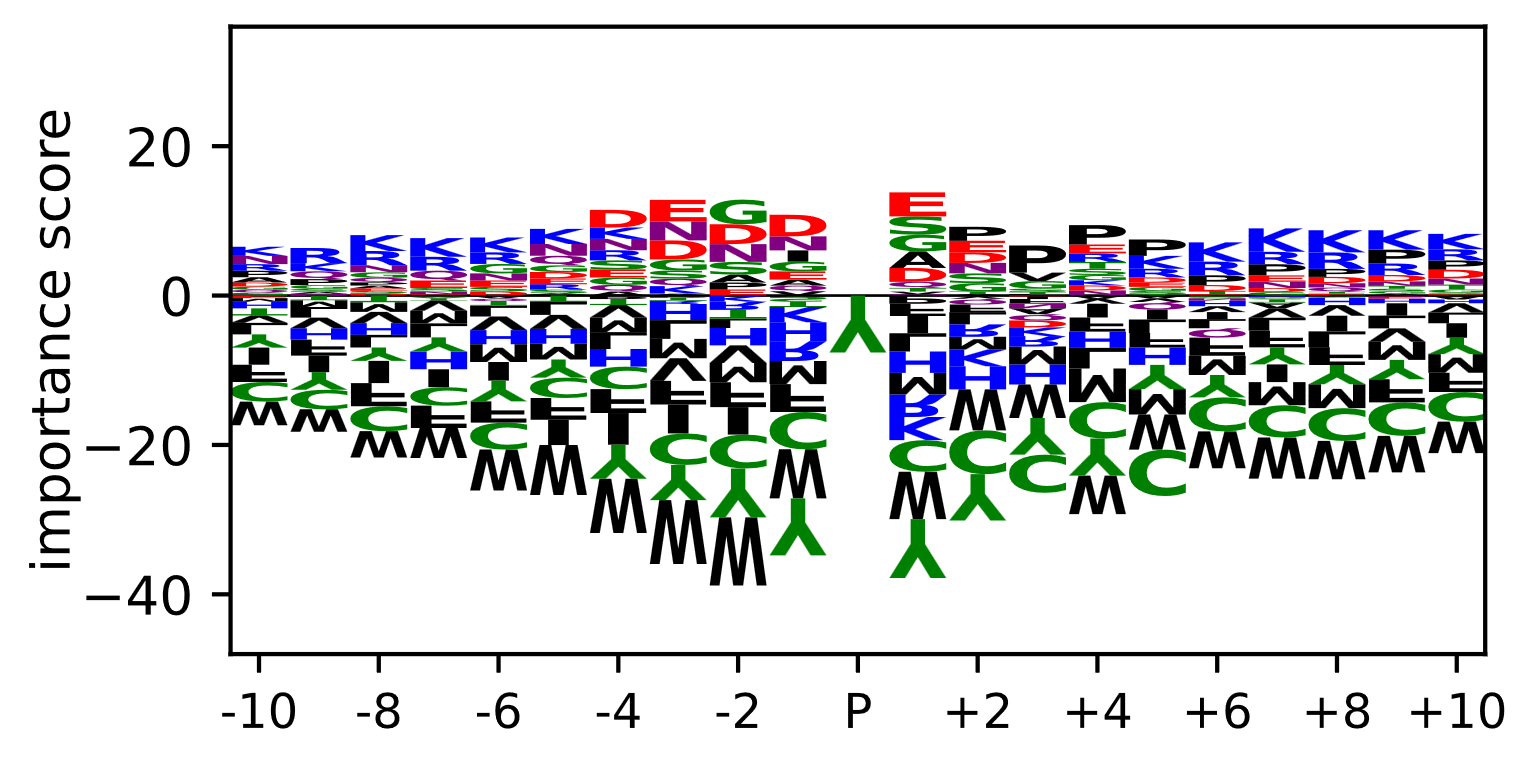 |

| ProtT5-XL-U50, trained on **Scop3P-ST-PF** data | ProtT5-XL-U50, trained on **Scop3P-Y-PF** data |
| --- | --- |
| 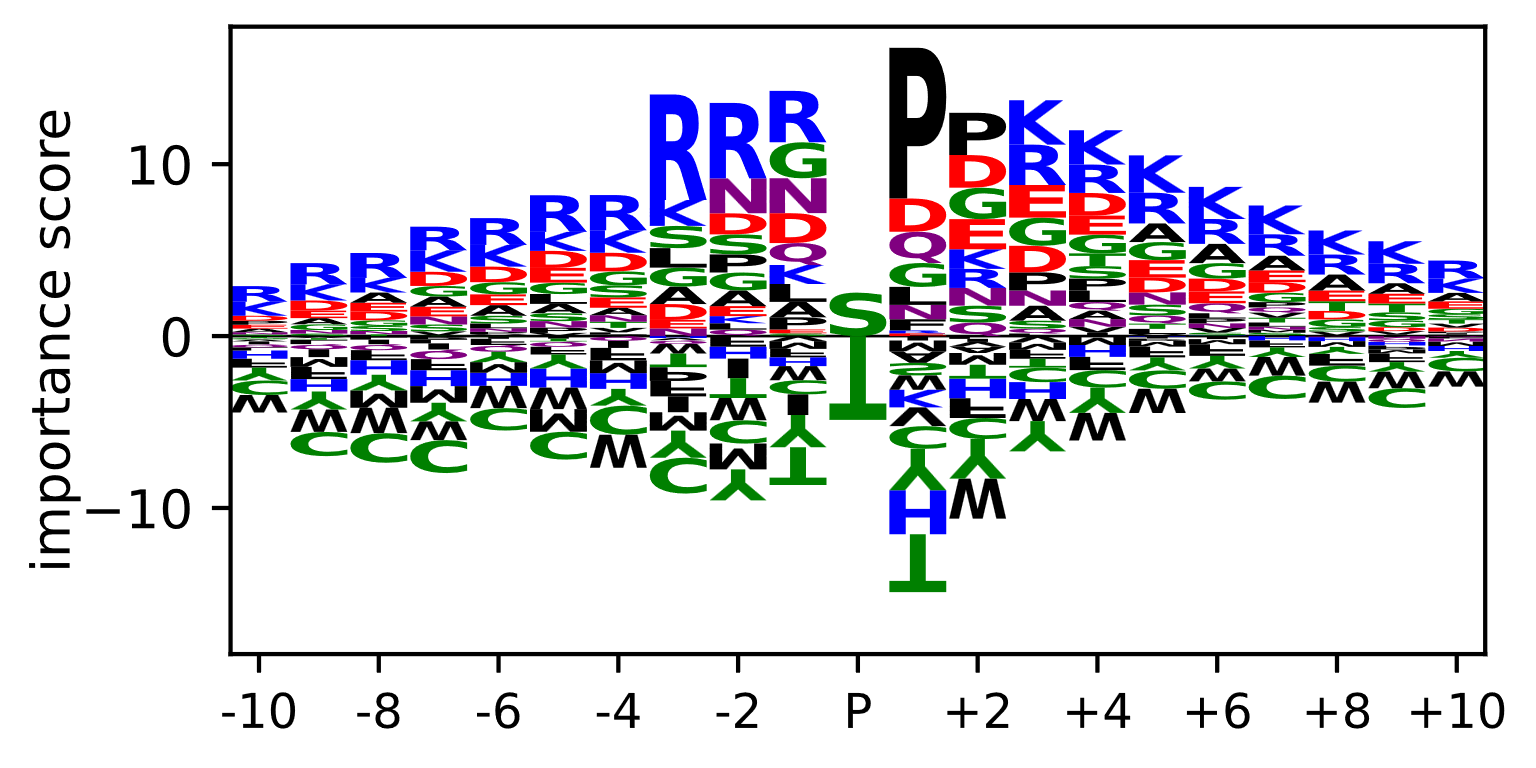 | 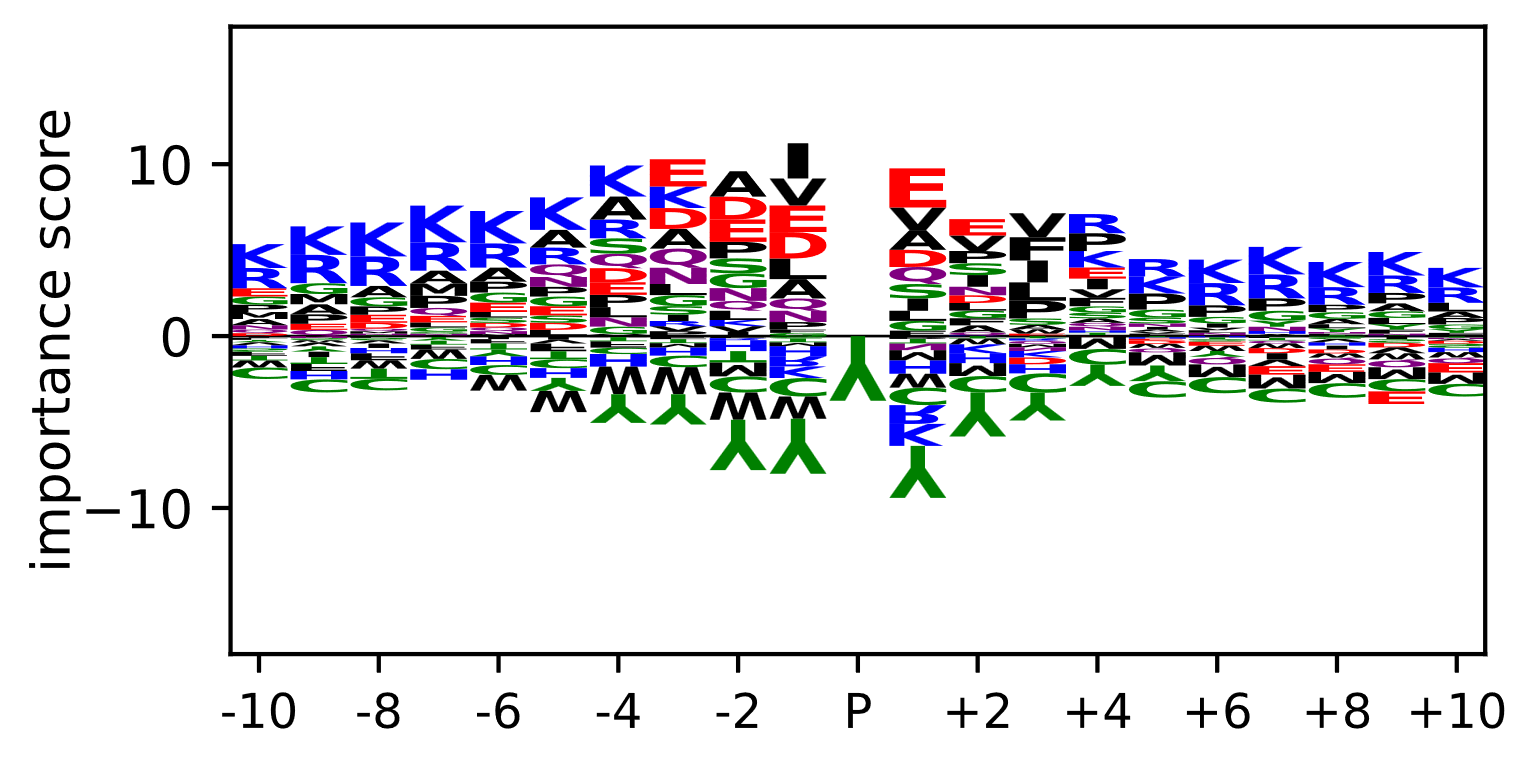 |

**Supplementary Figure 5.** Average importance scores per position, calculated using DeepLiftSHAP, for one-hot encoded and ProtT5-XL-U50 models trained on the peptide-filtered Scop3P datasets.

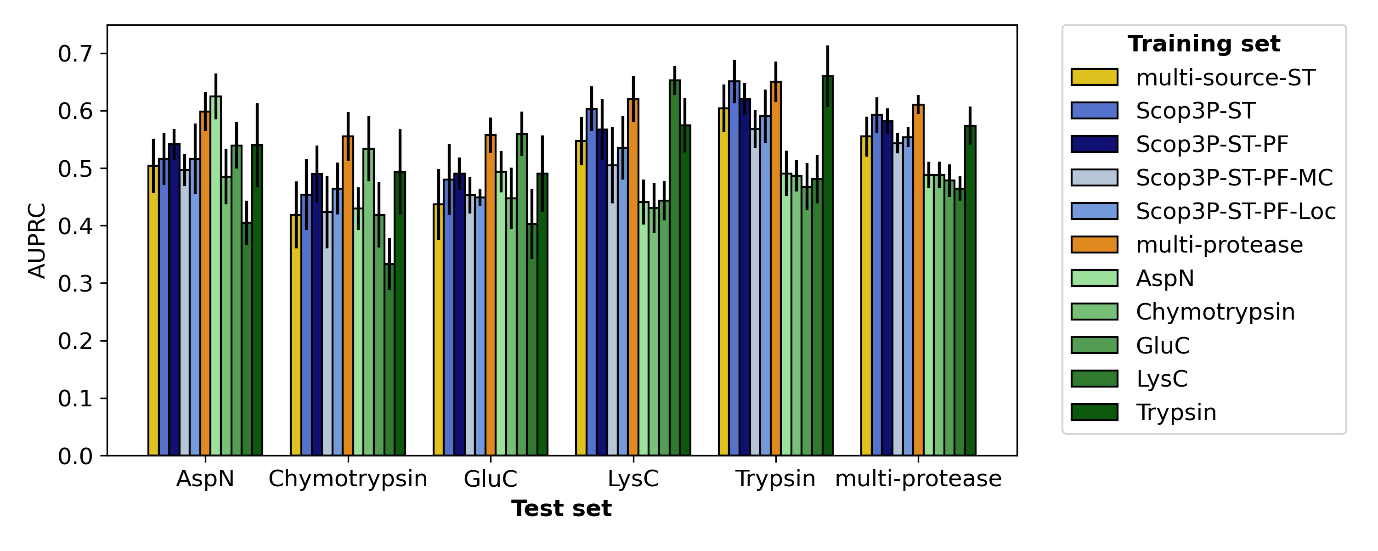
***Supplementary Figure 6.*** *Evaluation on non-peptide-filtered single-protease datasets, when training on another dataset first. The average AUPRC over ten test folds is shown, for the ProtT5-XL-U50 models. To avoid information leakage, proteins in the test fold were excluded from the training set.*

average localization score

| Peptide length | Series 1 |
| --- | --- |
| 5 | 1 |
| 6 | 0.887027 |
| 7 | 0.86758 |
| 8 | 0.858011 |
| 9 | 0.857945 |
| 10 | 0.831329 |
| 11 | 0.818555 |
| 12 | 0.809302 |
| 13 | 0.807999 |
| 14 | 0.793176 |
| 15 | 0.767084 |
| 16 | 0.758358 |
| 17 | 0.742157 |
| 18 | 0.705848 |
| 19 | 0.701131 |
| 20 | 0.709909 |
| 21 | 0.677578 |
| 22 | 0.671693 |
| 23 | 0.640822 |
| 24 | 0.626923 |
| 25 | 0.563425 |
| 26 | 0.605578 |
| 27 | 0.567609 |
| 28 | 0.567015 |
| 29 | 0.533683 |
| 30 | 0.543776 |
| 31 | 0.687146 |
| 32 | 0.635129 |
| 33 | 0.605347 |
| 34 | 0.524787 |
| 35 | 0.430934 |

***Supplementary Figure 7.*** *Average localization score per peptide length. Localization scores were calculated with PhosphoRS*^2^ *on all peptide-spectrum matches where exactly one phosphorylation was detected. The average localization scores for the phosphosites that were reported by the search engine (in this case, ionbot*^3^*) in Scop3P.*

| name | # of parameters | output size | availability |
| --- | --- | --- | --- |
| one-hot encoding | - | (n, 20) | - |
| ESM-1_small_ | 43M | (n, 768) | <https://github.com/facebookresearch/esm> |
| ESM-1b | 650M | (n, 1280) | <https://github.com/facebookresearch/esm> |
| ESM-2_150M_ | 150M | (n, 640) | <https://github.com/facebookresearch/esm> |
| ESM-2_650M_ | 650M | (n, 1280) | <https://github.com/facebookresearch/esm> |
| ESM-2_3B_ | 3B | (n, 2560) | <https://github.com/facebookresearch/esm> |
| CARP_640M_ | 640M | (n, 1280) | <https://github.com/microsoft/protein-sequence-models> |
| ProtT5-XL-U50 | 1.5B (inference) | (n, 1024) | <https://github.com/agemagician/ProtTrans> |
| Ankh_base | 450M | (n, 768) | <https://github.com/agemagician/Ankh> |
| Ankh_large | 1.15B | (n, 1536) | <https://github.com/agemagician/Ankh> |

**Supplementary Table 4**. Details for the different protein representations that were compared in this study.

| **hyperparameter** | **setting** |
| --- | --- |
| receptive field | 195 |
| # of conv blocks | 2 |
| # of conv filters | 325 |
| conv filter size | 15 |
| max pool size | 2 |
| # of neurons | 350 |
| dropout | 0.4 |
| learning rate | 1e-4 |
| batch size | 16 |
| optimizer | AdamW |
| # of warm-up epochs | 1.5 |
| early stopping | validation loss is calculated every 1/4^th^ epoch, and early stopping is done when loss does not decrease for 1.25 epochs. |
| max # of epochs | 15 |

**Supplementary Table 5.** The hyperparameters for the CNN and its training scheme as used throughout this publication. In an initial setup, a hyperparameter search was done on the Scop3P-ST dataset, for one-hot, ESM-1_small_, ESM-1b, and ProtT5-XL-U50 language models. As the number of available pLMs increased, we determined one hyperparameter setup by taking the consensus of those previous experiments.

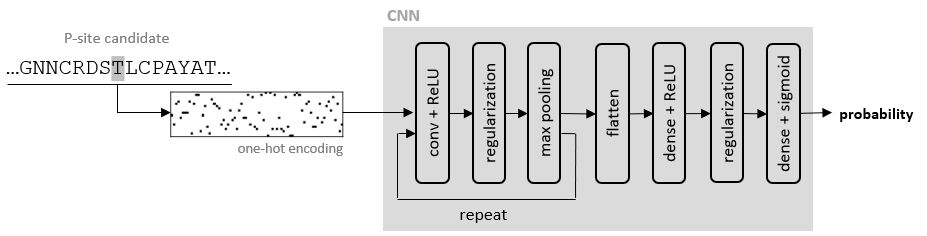

***Supplementary Figure 8.*** *One-hot encoded CNN: Convolutional neural network architecture for phosphorylation prediction, taking a one-hot encoding as input. The architecture starts with a series of convolutional blocks is, with each block consisting of a convolutional layer (with ReLU non-linearization), a regularization layer (either dropout or batch normalization), and a max pooling layer. The output of the last convolutional block is flattened and fed to a dense layer (with ReLU non-linearization), followed by a regularization layer, a dense layer with one output neuron, and a sigmoid.*

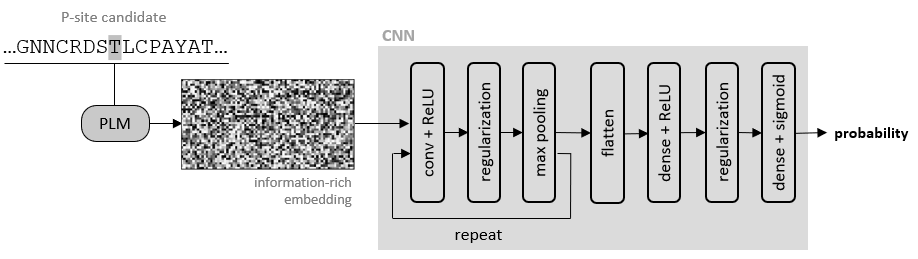

***Supplementary Figure 9.*** *The pLM-based setup uses the same architecture as the one-hot encoded models in Supplementary Figure 1, but now takes an information-rich protein representation computed by a pLM as input.*

| **name** | **source** | **residues** | **# of proteins** | **# of pos** | **# of neg** | **pos:neg ratio** |
| --- | --- | --- | --- | --- | --- | --- |
| multi-source-ST | Guo21^4^ | ST | 13,599 | 184,375 | 981,620 | 1:5.3 |
| multi-source-Y |  | Y | 9,710 | 32,213 | 149,501 | 1:4.6 |
| **name** | **source** | **residues** | **# of proteins** | **# of pos** | **# of neg** | **pos:neg ratio** |
| Scop3P-ST | Ramasamy22^5^ | ST | 8,754 | 54,395 | 681,975 | 1:12.5 |
| Scop3P-Y |  | Y | 8,754 | 4,884 | 125,306 | 1:25.7 |
| **name** | **source** | **protease** | **# of proteins** | **# of pos** | **# of neg** | **pos:neg ratio** |
| AspN | Giansanti15^6^ | AspN | 1,816 | 3,987 | 171,003 | 1:42.9 |
| Chymotrypsin | (ST only) | Chymotrypsin | 1,739 | 3,476 | 167,754 | 1:48.3 |
| GluC |  | GluC | 1,899 | 4,280 | 184,426 | 1:43.1 |
| LysC |  | LysC | 1,718 | 4,133 | 166,669 | 1:40.3 |
| Trypsin |  | Trypsin | 3,255 | 9,096 | 298,628 | 1:32.8 |
| multi-protease |  | multi-protease | 4,728 | 18,028 | 424,527 | 1:23.5 |

**Supplementary Table 6.** An overview of phosphorylation datasets used in this study.

| **name** | **source** | **residues** | **# of proteins** | **# of pos** | **# of neg** | **pos:neg ratio** |
| --- | --- | --- | --- | --- | --- | --- |
| Acetylation | PhosphoSitePlus^1^  + dbPTM^7^ | K | 15,737 | 64,029 | 666,016 | 1:10.4 |
| Methylation |  | KR | 7,493 | 18,824 | 705,204 | 1:37.5 |
| Sumoylation |  | K | 3,193 | 11,606 | 151,426 | 1:13.0 |
| Ubiquitination |  | K | 20,080 | 136,817 | 684,015 | 1:5.0 |

**Supplementary Table 7.** An overview of datasets for other PTMs. Due to the lack of metadata, no peptide filtering is applied here.

| **name** | **source** | **residues** | **# of proteins** | **# of pos** | **# of neg** | **pos:neg ratio** |
| --- | --- | --- | --- | --- | --- | --- |
| Scop3P -ST-PF | Ramasamy22^5^ | ST | 8,754 | 54,395 | 326,584 | 1:6.0 |
| Scop3P -ST-PF |  | Y | 8,754 | 4,884 | 60,468 | 1:12.4 |
| **name** | **source** | **protease** | **# of proteins** | **# of pos** | **# of neg** | **pos:neg ratio** |
| AspN-PF | Giansanti15^6^ | AspN | 2,182 | 3,987 | 5,475 | 1:1.4 |
| Chymotrypsin-PF | (ST only) | Chymotrypsin | 2,097 | 3,476 | 5,199 | 1:1.5 |
| GluC-PF |  | GluC | 2,374 | 4,280 | 6,280 | 1:1.5 |
| LysC-PF |  | LysC | 2,389 | 4,133 | 5,106 | 1:1.2 |
| Trypsin-PF |  | Trypsin | 3,923 | 9,096 | 9,479 | 1:1.0 |
| multi-protease-PF |  | multi-protease | 5,867 | 18,028 | 25,341 | 1:1.4 |

**Supplementary Table 8.** An overview of peptide-filtered phosphorylation datasets considered in this study. Note that the filtering step was not done for the multi-source data due to the lack of metadata.

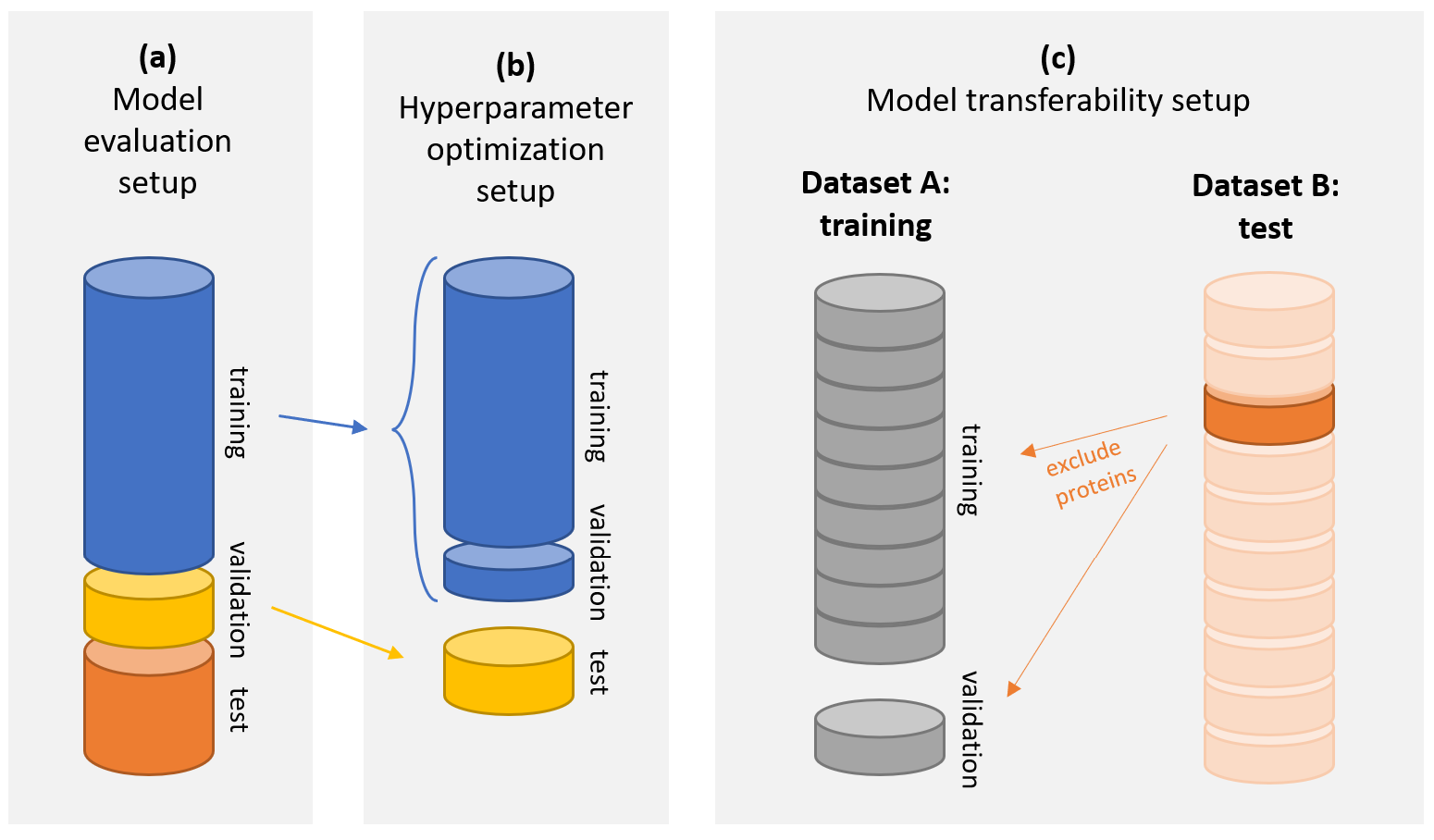

**Supplementary Figure 10.** An illustration of the dataset splits during the different experimental setups: (a) fixed training, validation and test sets are used for the comparison of different predictors, (b) the training and validation sets of the fixed splits were used for the initial hyperparameter search that helped determining the final settings, and (c) in the model transferability setup, models are evaluated on a test set, divided into ten folds, and training is done on another dataset from which all proteins in the test fold are excluded to avoid data leakage.

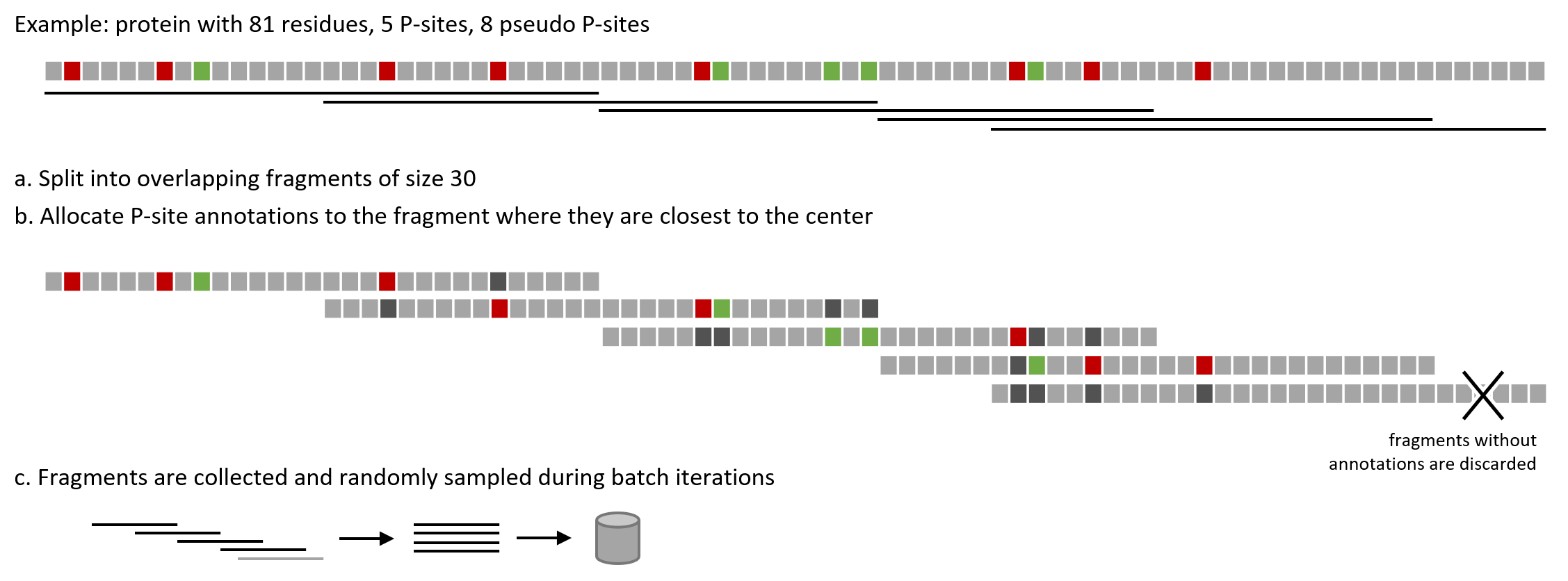

**Supplementary Figure 11**. An illustration of how sequences are effectively split into overlapping fragments, and how phosphosite annotations are distributed among them (example with fragment length 30 instead of 512, for the sake of clarity). (a) The protein is split into overlapping fragments, with an overlapping size of half the sequence. The last fragment will have its starting position shifted to the left in order to maintain the same sequence length as the other fragments. (b) phosphosite annotations are distributed among fragments, where they are tied to the fragment where they are closest to the center. (c) Fragments from the same protein can be present in different batches. However, when splitting datasets into training, validation, and test, fragments of the same protein are found in the same set.
